# Analysis of translating mitoribosome reveals functional characteristics of translation in mitochondria of fungi

**DOI:** 10.1101/2020.01.31.929331

**Authors:** Yuzuru Itoh, Andreas Naschberger, Narges Mortezaei, Johannes M. Herrmann, Alexey Amunts

## Abstract

Mitoribosomes are specialized protein synthesis machineries in mitochondria. However, how mRNA binds to its dedicated channel, and tRNA moves as the mitoribosomal subunit rotate with respect to each other is not understood. We report models of the translating fungal mitoribosome with mRNA, tRNA and nascent polypeptide, as well as an assembly intermediate. Nicotinamide adenine dinucleotide (NAD) is found in the central protuberance of the large subunit, and the ATPase inhibitory factor 1 (IF_1_) in the small subunit. The models of the active mitoribosome explain how mRNA binds through a dedicated protein platform on the small subunit, tRNA is translocated with the help of the protein mL108, bridging it with L1 stalk on the large subunit, and nascent polypeptide paths through a newly shaped exit tunnel involving a series of structural rearrangements. An assembly intermediate is modeled with the maturation factor Atp25, providing insight in to the biogenesis of the mitoribosomal large subunit and translation regulation.

## Introduction

Protein synthesis in mitochondria supports bioenergetics of eukaryotic cells and is executed by dedicated mitoribosomes. Electron cryo-microscopy (cryo-EM) has been instrumental in revealing the first structural snapshots of mitoribosomes. While for mammals the obtained models from human embryonic kidney cells ^1,2^ are in agreement with those from porcine and bovine tissues ^3,4^, the structure of the mitoribosome from the yeast *Saccharomyces cerevisiae* displayed considerable specializations ^5,6^. Particularly, it proposed a distinct composition and putative deviations in the functionally defining features of translation, including an expanded mRNA channel exit and rerouted polypeptide exit channel. Although the existing cryo-EM data is informative, the deviations of the mitoribosome from *S. cerevisiae* that lacks complex I are more likely species-specific and therefore cannot be considered as a prototypic example of fungi. In addition, the available models are incomplete due to the limited resolution. Therefore, in the absence of high-resolution structural information of the translating mitoribosome, key questions regarding mitochondrial translation remain open. To provide a representative reference for studying protein synthesis in the mitochondria of fungi, and to reveal how the mitoribosome functions in coordination with its translation partners, we determined structures of the translating mitoribosome from the representative fungal model organism *Neurospora crassa*.

*N. crassa* has been recognized as the organism used for answering a variety of fundamental biological questions in the field of mitochondrial translation, and low resolution reconstruction of the mitoribosome was reported ^7^. Other contributions of this model organism include early characterization of the physicochemical properties of the mitoribosomes and the base composition of its rRNA ^8^, the finding that mitoribosomal intactness requires a relatively high magnesium concentration ^9^, studies showing radioactive amino acid incorporation into mitoribosome ^10^, and the finding that Oxa1 facilitates integration of mitochondrially encoded proteins into the inner membrane ^11^. Therefore, to provide a reference for the process of protein synthesis in mitochondria, we set out to investigate the functional translation apparatus from the model organism *N. crassa*.

The work shows that mitoribosomes acquire cofactors and subunits associated with the respiratory complexes, such as NAD and IF_1_. Binding of mRNA requires extended mitoribosomal proteins of the small subunit, and movement of tRNA is realized through additional proteins of the large subunit. Evolutionary analysis comparing mitoribosomes from different species with bacterial counterparts, illustrates that the exit tunnel evolves via deletions in the rRNA and extensions of mitoribosomal proteins. Finally, we describe a bL9m-lacking assembly intermediate complexed with the maturation factor Atp25 that is formed as a result of protein splitting.

## Results and Discussion

### Structure determination

In order to characterize a representative functional mitoribosome, the *N. crassa* mycelia of the strain K5-15-23-1 overexpressing the protein Oxa1 ^11^ were grown aerobically and translation was stalled using chloramphenicol prior to harvesting. The mycelia were dried, lysed with mortar and pestle, and the mitoribosomes were purified (see Methods), and subjected to cryo-EM analysis. The cryo-EM consensus map was obtained from 131,806 particles at overall resolution of 2.83 Å (Supplementary Fig. 1 and Supplementary Table 1).

The 3D classification resulted in two reconstructions of the mitoribosome corresponding to the rotated and non-rotated states (Supplementary Fig. 1). After subtracting the signals outside the tRNA binding sites, followed by focused 3D classifications, we obtained three distinct classes with tRNAs occupying P-site (P/P), E-site (E/E), and in the hybrid P/E state, at the overall resolutions of 3.05, 3.10, and 2.99 Å, respectively (Supplementary Fig. 1). In the P/P and P/E tRNA states, we detected and modeled co-purified native mitochondrial mRNA, and the density corresponding to the nascent polypeptide was identified in the P/P tRNA state.

To further improve the local resolution, focused masked refinements were performed using the masks for the large subunit (mtLSU) core, central protuberance (CP), L1 stalk, uL10m region, the small subunit (mtSSU) head, body and tail (Supplementary Fig. 2–5). It allowed identification of five proteins previously missed in the yeast structure: uL1m, uL10m, uL11m, mL53, and mL54; and two additional mitoribosomal proteins: mL108 and the ATP synthase inhibitory factor 1 (IF_1_); and 60 proteins with higher completeness, as well as the rRNAs (Fig. 1a, Supplementary Fig. 6–10).

**Fig. 1:**
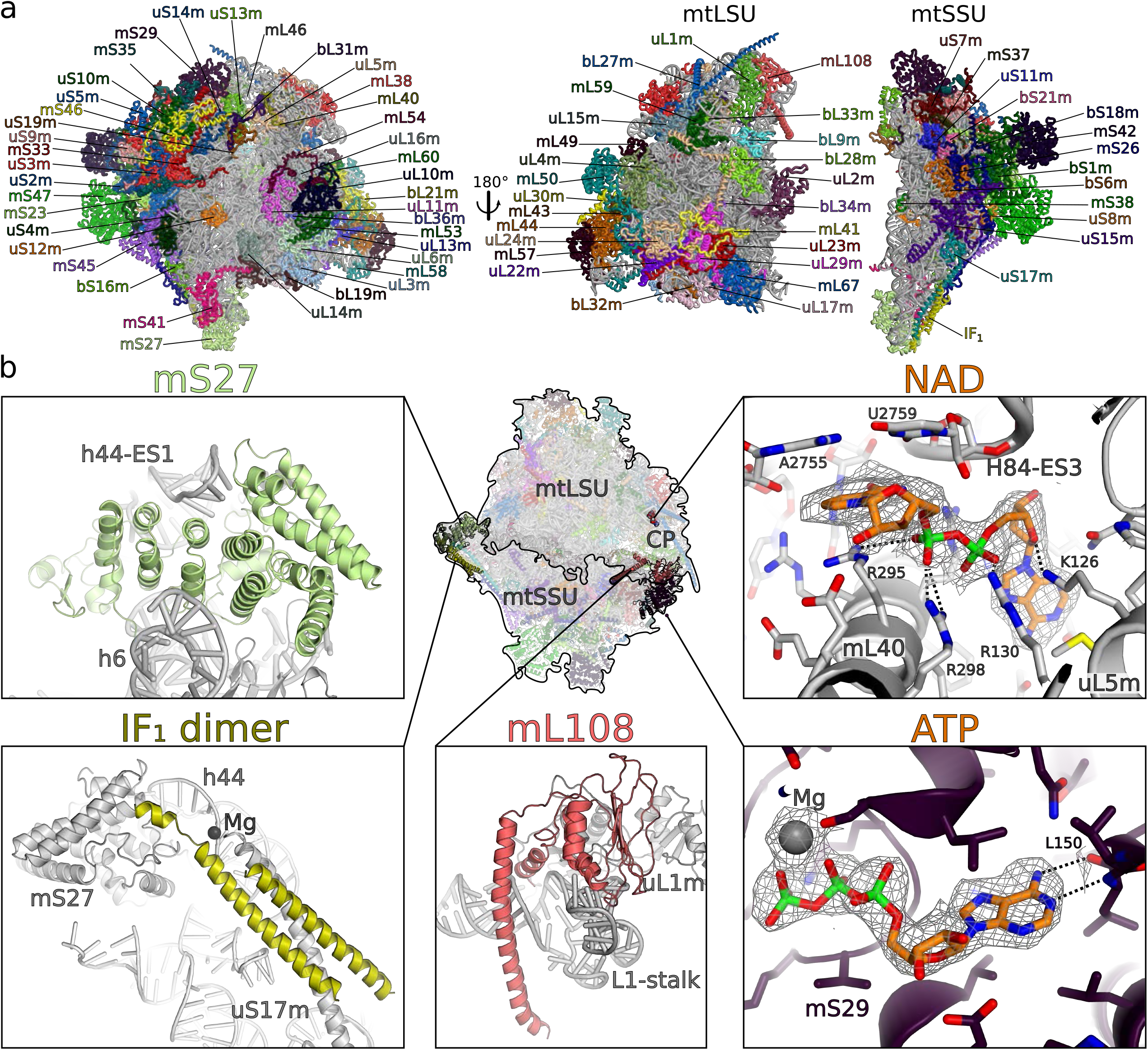
Structure of the fungal mitoribosome and additional features. **a** Overall structure of the complete mitoribosome and its subunits with indicated mitoribosomal proteins. **b** Examples of additional features. Remodeled and reannotated mS27 interacts with h44-ES1. The mitoribosomal component IF_1_ dimer bound to h44-ES1 and uS17m extension. NAD binding pocket in the CP formed by uL5m, mL40, and H84-ES3. The mitoribosomal component mL108 is located in the L1 stalk. Nucleotide binding pocket of mS29 bound with chemically favorable ATP shown with its density.

We also noted that mitoribosomes from actively growing *N. crassa* mycelia contain a pool of free mtLSU (Supplementary Fig. 5), which resembles the profile of the assembly intermediate reported from the mitochondria of human embryonic kidney cells ^12^. Unlike the mtLSU in the intact mitoribosome, this population is missing the density for bL9m, but has an extra factor Atp25 bound to uL14m, and a poor density at the interface consistent with unfolded mt-rRNA. The quality of the structural data allowed the analysis of the differences with the mature mtLSU that suggests a putative regulatory mechanism.

### Overall structure and additional features

The fungal mitoribosome consists of 78 different proteins, 23S and 16S rRNAs (Fig. 1, Supplementary Fig. 6–10, Supplementary Tables 2 and 3). The proteins mS42 and IF_1_ are found in two copies as homodimers. 48 bacterial orthologs are extended, and 30 proteins are mitochondria-specific. For the rRNA, we modeled 16 expansion segments (ES), 15 of which are in the mtLSU (Supplementary Fig. 7–9), and several other rRNA regions have been reduced. The deletion in the mtDNA genome are not correlated with a specific functional feature or attributed to selective loss/gain of activity, but rather reflect a general reductive phase in the mitochondrial genome ^7^. For the native mitochondrial tRNAs (P/P, P/E, E/E states) and mRNA, we modeled nucleotide sequences that fit the corresponding densities the best, because those densities represent a mixture of different tRNAs and mRNAs in mitochondria.

Some of the improvements in the model due to the high-resolution map are exemplified in Fig. 1. Protein mS27, which was previously partially built as poly-Ala, has been identified directly from the density as a helical repeat protein consistent with mammalian counterparts. We named it accordingly mS27 (previously mS44) (Fig. 1b, Supplementary Fig. 11, Supplementary Table 3). Another example is the protein mS29 that was reported as a GTPase involved in apoptosis, and mutations in the P-loop motif of the nucleotide binding pocket were suggested to impair this function ^13^. In the nucleotide pocket of mS29, which was previously modeled with guanosine diphosphate (GDP), we can see at the local resolution of 2.77 Å that the density for N2 purine ring atom is missing, and the correct assignment for the nucleotide is adenosine triphosphate (ATP) (Fig. 1b, Supplementary Fig. 12). This is further supported by the better resolved chemical environment formed by Leu150 of mS29, which would be incompatible with guanine since the O6 carbonyl of guanine imparts repulsive interaction with the backbone carbonyl of Leu150 (acceptor with acceptor), and the NH1 group of guanine forms a second repulsive interaction with the backbone NH group of Leu150 (donor with donor). Placing an ATP instead of GDP leads to more favorable interactions.

In the CP, we found an extra density in the pocket formed by mL40, uL5m, and rRNA H84-ES3 (Fig. 1b). Based on the density, interactions and the abundance in mitochondria, we conclude that the it is most likely to be NAD (Supplementary Fig. 13). Positively charged residues of mL40 and uL5m interact with the negatively charged NAD phosphates. The pyridine ring of the nicotinamide is held in place by π-stacking on the base pair of A2755 and U2759. Arg295 of mL40 forms π-stacking at the other side of the NAD pyridine. Therefore, NAD bridges the mitoribosomal proteins and the mt-rRNA at the core of CP. This implies regulatory function in the assembly of the mtLSU based on the local NAD level.

In the mtSSU tail, we found an extra density corresponding to helical structures of protein chains. The density (Supplementary Fig. 6 and 14) and the mass-spectrometry data (Supplementary Table 4) suggested that it is the natural inhibitor of the mitochondrial ATP synthase IF_1_ bound as a homodimer (Fig. 1b). In the ATP synthase, IF_1_ functions through inserting its N-terminal part into the catalytically active F_1_-ATPase, thereby blocking its rotational movement and subsequently the ATP hydrolase activity ^14,15^. In our structure of the mitoribosome, IF_1_ coiled-coil forms a helical bundle with the C-terminal extension of uS17m and also binds to mS27 and h44 rRNA (Fig. 1b, Supplementary Fig. 14). It is also bridged through a metal ion Mg^2+^, coordinated by uS17m Q162 and the backbone phosphate of h44 A1745 (Supplementary Fig. 2c). The topology of IF_1_ homodimer is such that the C-terminus of each monomer points in the opposite direction: one towards the mtSSU body, and the other towards the solvent. Although the topology resembles that of the bovine IF_1_ homodimer ^16^ (Supplementary Fig. 14a), the dimerization region is shifted ~7 residues toward the N-terminus and is shorter by ~10 residues, suggesting it is less stable. In addition, against the sequence similarity with the yeast counterpart (Supplementary Fig. 14b), we find differences in topology when compared with the X-ray crystal structure of the *S. cerevisiae* IF_1_ monomer bound to the F_1_-ATPase ^14^ (Supplementary Fig. 14a). Particularly, the *S. cerevisiae* IF_1_ has a short helix α1 followed by a longer helix α2, whereas in *N. crassa* the α1 counterpart region appears to be disordered, and α2 comprises two helices separated by 2 residues. The differences might also be affected by different binding partners (mitoribosome vs F_1_-ATPase), reflecting the structural flexibility of IF_1_. Since the C-terminal extension of uS17m stabilizing the IF_1_ on the mitoribosome is specific to *N. crassa* (Supplementary Fig. 14c) the feature of IF_1_ binding might also be specific. Our structural analysis shows that h44 is expanded in this region of the mitoribosome, which would suggest that IF_1_ has been acquired through the mechanism of ‘structural patching’ to stabilize the rapidly evolving growth ^17^.

### The mechanism of mRNA binding involves a dedicated protein platform

Focused classifications on the tRNA binding sites yielded structures of three distinct classes with bound tRNA in P/P, P/E, E/E (Supplementary Fig. 1). The quality of the density allowed us to generate structural models of the mitoribosomal complexes with P/P and P/E tRNAs. The presence of tRNAs correlates with additional density lining the mRNA channel path, where a well resolved map allowed modeling of 11 nucleotides of the native mRNA (see Methods). The codon-anticodon base pairings are observed in the density of the P/P and P/E states (Fig. 2). The decoding center is conserved, including the position of the bases A1803 (A1492 in *E. coli*), A1804 (A1493) and G756 (G530), as well as the curvature in mRNA around h28. In addition to the modeled mRNA nucleotides in the channel, bulky density extends upstream to the channel entry and downstream to its exit. Therefore, the map allows us to determine how mRNA binds to the mitoribosome and to trace its complete path.

**Fig. 2:**
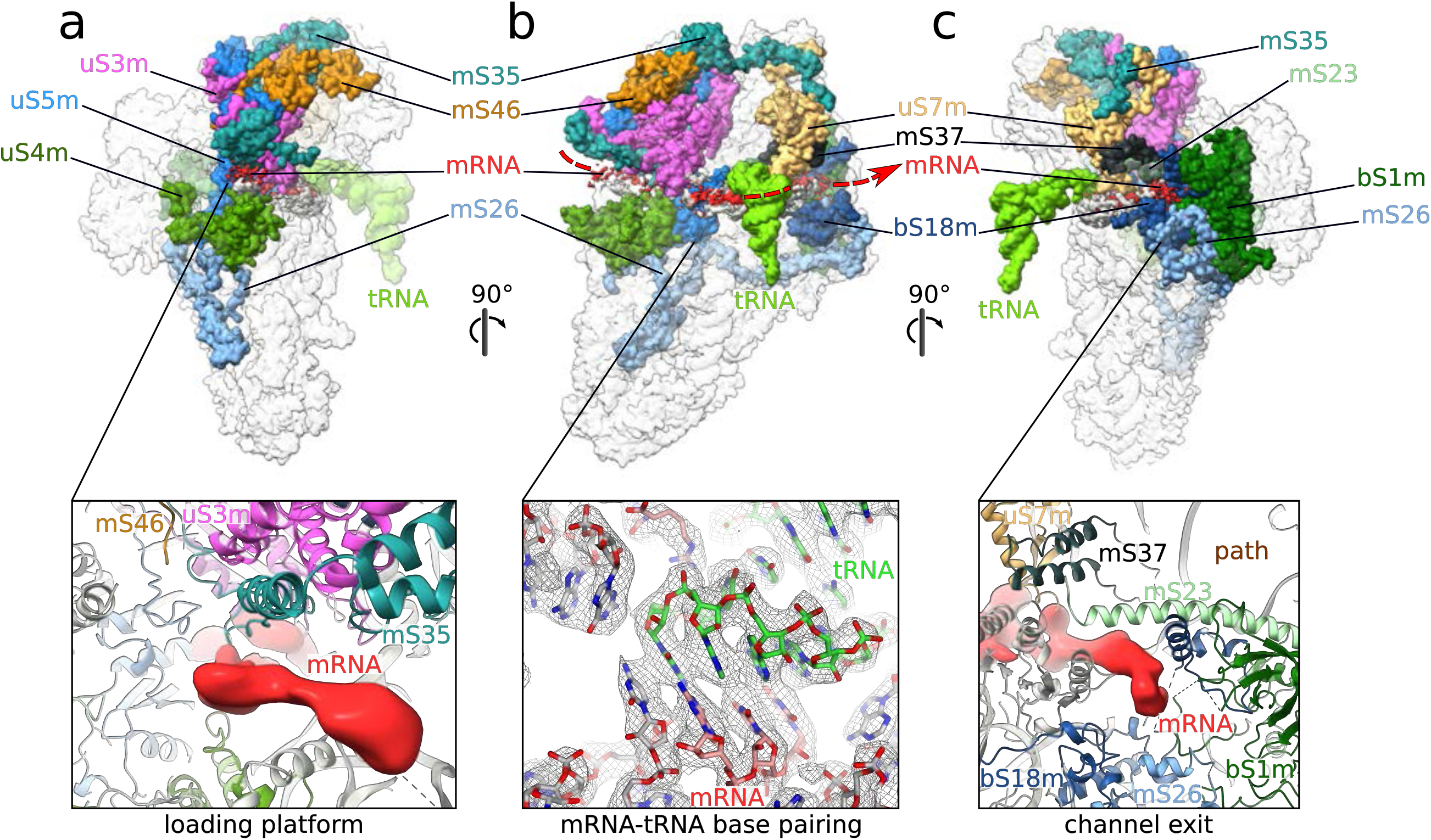
The mRNA channel traced with the density. **a** The density for mRNA (red) reveals a protein platform formed by mS35, mS46 and extensions of uS3m and uS5m. **b** The overall path taken by mRNA is indicated based on the additional density associated with the mtSSU, and codon-anticodon base pairings at the decoding center shown with density. (C) At the exit site, proteins uS7m, bS18m, mS23, mS26, and mS37 are in contact with mRNA, and bS1m is permanently bound. A putative alternative path is indicated.

The reference point to the channel entry is the universal site formed by uS3m, uS4m, and uS5m. Compared to the previously determined structures of the bacterial complex, mitochondrial mRNA is also bound by mitochondria-specific protein mS35. However, the density for the mRNA chain starts prior to the conventional entry. Particularly, following the density, we observe extended structure involving mitochondria-specific extensions of uS3m and uS5m that are held together at the mtSSU head by mS35 and mS46 (Fig. 2a). This architecture is coupled to the mRNA terminus, thereby suggesting a potential platform for its loading on to the mitoribosome. The formed loading platform narrows the entry site considerably, ensuring that the mRNA entering the channel is unpaired. Interestingly, two mitochondria-specific proteins, mS26 and mS35, extend along the mRNA channel from its entry region to the opposite mtSSU pole, where the exit site is (Fig. 2b).

The channel exit site typically resides at the 3’ end of the rRNA. In the fungal mitoribosome, it has not been significantly altered. The remodeling reported in *S. cerevisiae* does not occur, and therefore represents either a dormant state or species-specific feature. In addition to the conserved proteins, mitochondria-specific mS23, mS26, and mS37 shape the path for mRNA movement (Fig. 2c). The C-terminal extension of uS7m narrows the channel, and protein bS18m, mS23, mS26, and mS37 interact with the mRNA density directly. The path toward the exit appears to bifurcate into two subways, and each could in principle accommodate a single stranded mRNA. However, the density for the 5’ mRNA clearly only follows the path lateral to mS26 that is also surrounded by a positively charged chemical environment. Protein bS1m, which is considered to have one of the most conserved and ancient protein domains with functional importance of unfolding mRNAs for active translation ^18^, is permanently bound to the channel exit (Fig. 2c).

### The mechanism of tRNA translocation involves additional protein in L1 stalk

During translation, tRNAs are translocated between the three sites. Upon peptide-bond formation, the ribosomal subunits rotate with respect to each other, and the tRNA moves to the intermediate hybrid state. The structure comparison between the complexes with P/P and P/E tRNAs reveals sequential rearrangements in the network of mitoribosome-tRNA interactions and allows us to determine how tRNA is moved.

In the P/P state, the anticodon of the tRNA forms base pairing with the mRNA in the conserved P-site of the mtSSU (Fig. 3 and Supplementary Fig. 15). Further stabilization is provided by the conserved C-terminal Arg315 of uS9m, which forms salt bridges with the backbone phosphates in the anticodon loop of the tRNA. The conserved H69 of mtLSU interacts with the D-arm of the tRNA. The conformation of P/P tRNA is overall similar to bacteria. The A-site finger, known as a functional attenuator and important for keeping the correct reading frame ^19^, takes in a straight conformation forming the inter-subunit bridge B1a. The conserved G2453 (G2251) and G2454 (G2252) of the P-loop form base pairs with the CCA terminus of the tRNA, and the emerging peptide is bound to the terminal A76, waiting for peptidyl transfer to the next incoming aminoacyl-tRNA at the A-site. Thus, the arrangement of structural elements and tRNA in mitochondria of fungi shows a conserved P-site.

**Fig. 3:**
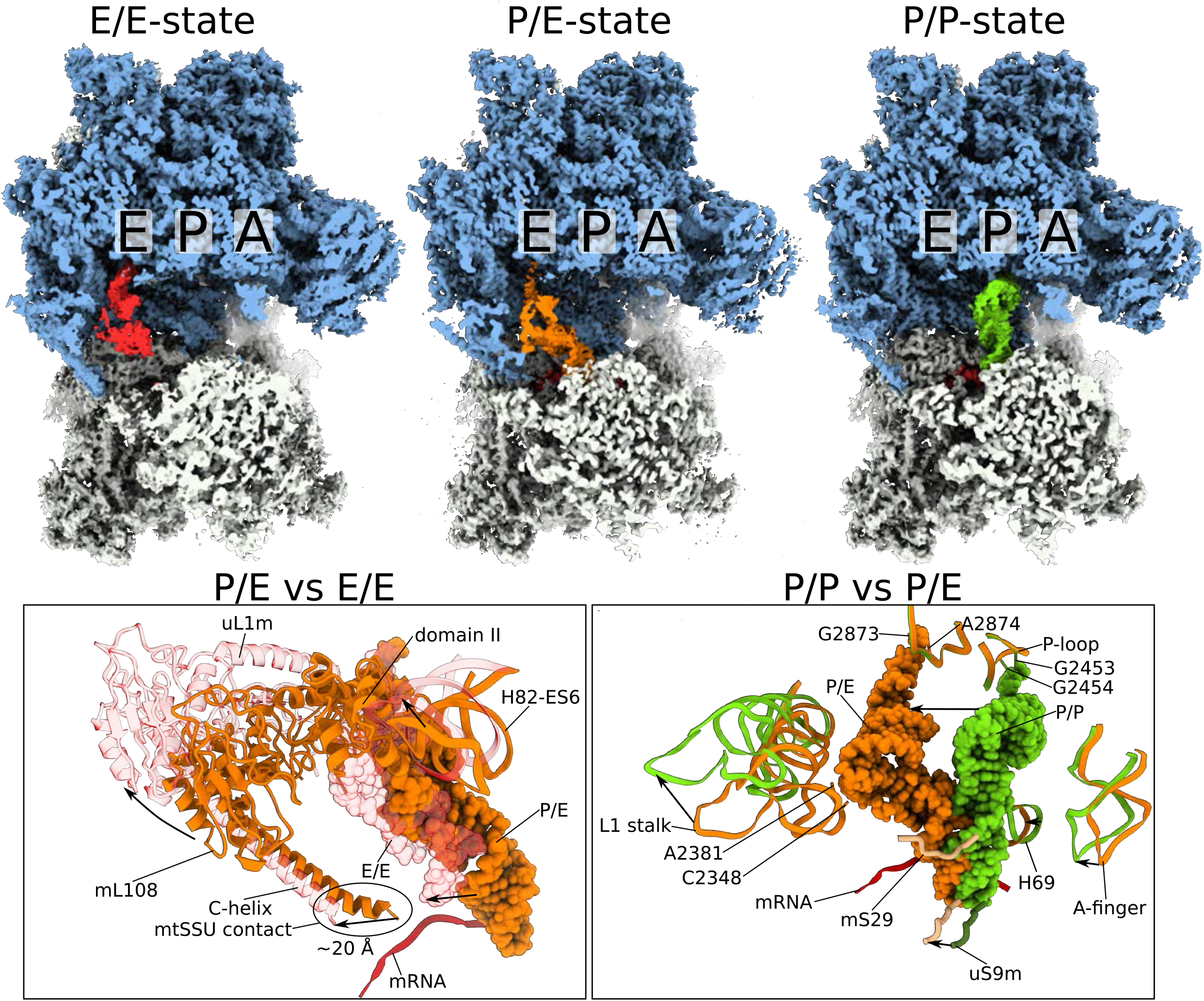
Translocation of tRNA and the L1 stalk. 3D reconstructions of the different states are shown for the E/E tRNA state for the P/E tRNA state and for the P/P tRNA state (mtLSU blue; mtSSU gray). Structural rearrangements upon transition between the P/E to E/E states and € between the P/P and P/E state are shown. Each two states have been superimposed, and the affected structural elements are shown according to the text. Movements of the structural elements between the states are indicated by arrows and the contact site to mtSSU is encircled.

After peptide transfer occurs, the ribosomal subunits move relative to each other, which is represented by our P/E hybrid structure. In the rotated conformation, the L1 stalk moves to a closed conformation toward the deacylated tRNA in the P/E hybrid state (Fig. 3). By masked refinement on L1 stalk, we improved the density that allowed building its complete model. We identified an additional protein component, namely mL108 (Fig. 1, Supplementary Fig. 6). The core of the protein shows a thioredoxin-like fold, which belongs to a small folding family of mitochondrial proteins mS25, mL43, and the complex I subunit NDUFA2/B8, bearing the possibility of a shared evolutional origin (Supplementary Fig. 16). The protein mL108 interacts with both the L1-stalk rRNA and the protein uL1m (Fig. 3). Furthermore, it forms an additional inter-subunit bridge with a long C-terminal helix (Fig. 3).

The bases A2381 (G2168) and C2348 (G2112) of the stalk rRNA stack on the elbow of the P/E tRNA (Fig. 3). Unlike in bacteria, uL1m does not interact with the P/E tRNA, instead the domain II of uL1m changes its conformation to form a mitoribsome-specific interaction with H82-ES6 in the CP in the rotated state (Fig. 3). Therefore, the subunit rotation is synchronized with the L1 stalk movement during tRNA translocation. The terminal adenosine of the P/E tRNA inserts between the conserved G2873 (G2421) and A2874 (C2422) at the E site (Fig. 3). The anticodon arm of the tRNA remains located in the P-site forming the codon-anticodon base pairing with the mRNA resulting in distorted conformation to swing the acceptor arm into the E-site. The disordered N-terminal extension of mS29 gets structured in the rotated state and interacts with the minor groove of the anticodon stem of the P/E tRNA (Supplementary Fig. 15), suggesting mitochondria-specific regulation. The long C-terminal helix of mL108 reaches down to the mtSSU, applying an additional stabilization of the L1 stalk.

Upon back rotation of the mtSSU into the non-rotated E/E state, the L1 stalk moves outwards assisting to pull the P/E tRNA into the E/E state, allowing dissociation into the solvent to complete the translocation cycle (Fig. 3 and Supplementary Fig. 15). The protein uL1m thereby detaches from the CP, releasing the domain II of uL1m. The tip of the C-terminal helix of mL108 translates ~20 Å forming a bridge with mtSSU. This structural rearrangement during mtSSU rotation suggests a coordinated movement of the mtSSU rotation and mL108 to the L1 stalk. The involvement of mitochondria-specific elements in coordination of tRNA movement has been recently reported also in other systems ^20–22^.

Taken together, the data enables to describe the tRNA movement on the mitoribosome and a series of related conformational changes. First, the L1 stalk interacts with mtSSU through mL108 thereby sensing the rotation state. This leads to the inward moving of the rRNA part of the L1 stalk to interact with the tRNA. The domain II of uL1m stabilizes the conformation through the mitochondria-specific interactions with the CP, and mS29 N-terminal loop becomes ordered and stabilizes the tRNA in the P/E state, which is disrupted in the following E/E state.

### The nascent polypeptide path to exit the mitoribosome is driven by mitochondria-encoded rRNA reorganisation

As the small mitoribosomal subunit progresses toward the 3’ end of the mRNA, the peptidyl transferase center transfers amino acids from tRNAs to the nascent chain, which emerges through the exit tunnel of the large subunit. Therefore, the polypeptide exit tunnel is the defining feature of protein synthesis, and it further facilitates initial folding. Given the fundamental role, the architecture of the tunnel path is among the most conserved features of the ribosome ^23^. Against this background, the *S. cerevisiae* mitoribosome structure showed a tunnel path that deviates from the bacterial counterpart, primarily due to rRNA H24 deletion ^5^. Consequently, the original tunnel was blocked by a specific extension of the mitoribosomal protein uL23m. To what extent the reported structural changes around the mitoribosomal exit tunnel are representative remained an open question.

In our structure, the density for an endogenous polypeptide is detected and the involved structural elements at the upper part of the tunnel are conserved with respect to the geometrical parameters, such as the length (~30 Å) and average radius (~5.3 Å) (Supplementary Fig. 17a). The constriction site between the loops uL4m-uL22m is located 34 Å from the start of the tunnel, and the position correlates well with the *E. coli* ribosome. The density for the nascent polypeptide can be traced from the starting point at the peptidyl transferase center (PTC) until the beginning of the lower part of the tunnel (Fig. 4a).

**Fig. 4:**
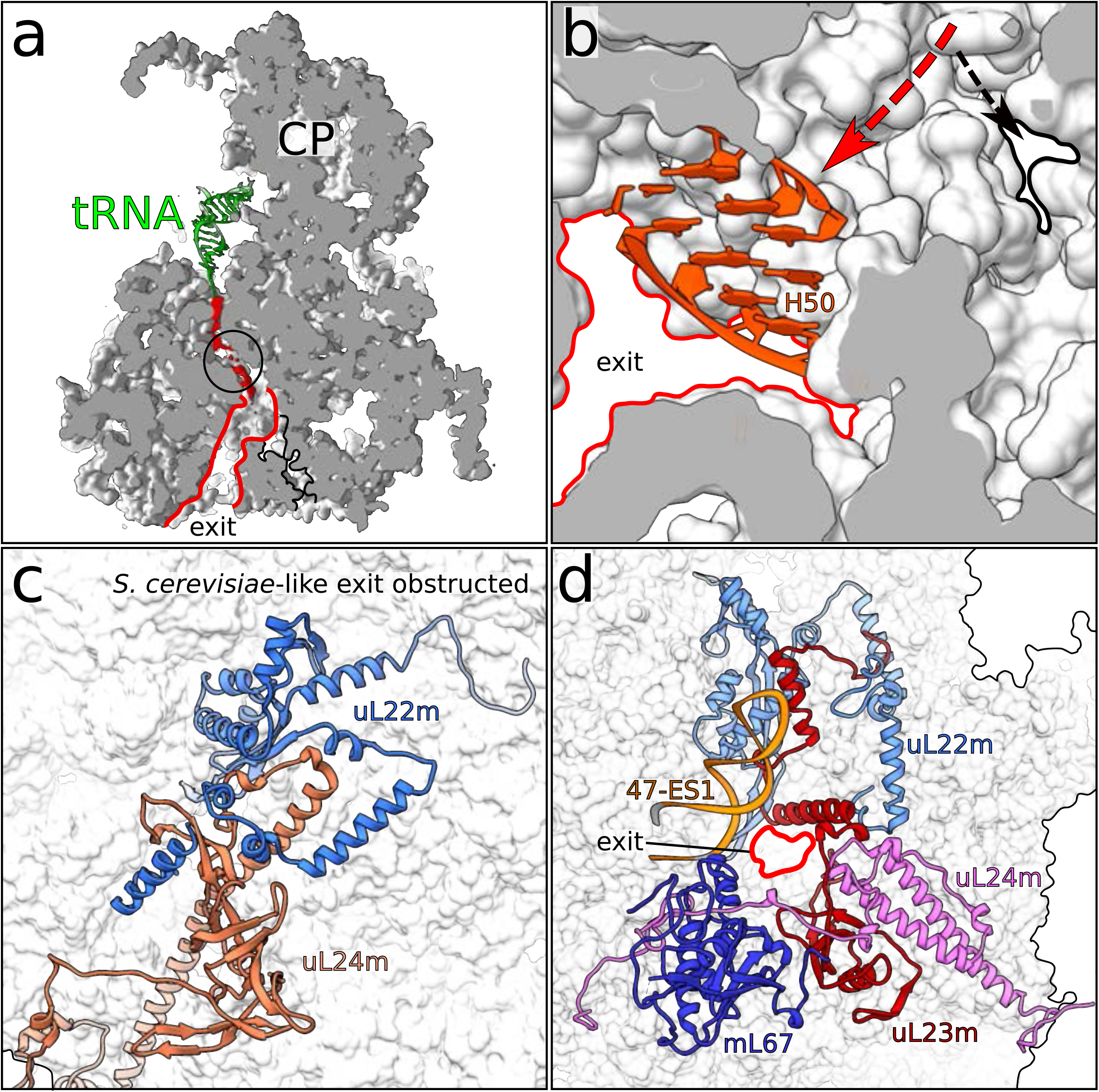
The path of polypeptide of the mitoribosome from *N. crassa*. **a** Cross-section of the large subunit showing the tunnel with density of the polypeptide reaching from the PTC until the beginning of the lower part of the tunnel. The constriction site is shown by a black circle and the constricted tunnel is highlighted by a black outline. **b** View of the branching point from inside the tunnel. The red arrow shows the exit which was formed upon deletion of H50 (orange). The black arrow shows the path of the constricted tunnel. **c** *S. cerevisiae*-like exit site is sterically hindered by protein extensions of uL22m and uL24m, shown in cartoon. **d** The view of the exit site showing the surrounding proteins and rRNA.

In the lower part, protein extensions (uL22m, uL23m, uL24m) cluster and occupy the conventional bacteria-like path interior. As a result, although a continuous channel can be traced to the surface of the mitoribosome, the minimal radius of the channel is decreased to 3.3 Å, which represents a considerable narrowing compared to ~6 Å across bacterial ribosomes ^23^ (Fig. 4a-b). We further find side chains that would obstruct the pathway, the majority of which belong to charged amino acids (Supplementary Fig. 17b). Such occupation would induce an electrostatic Coulomb potential that is inconsistent with hosting hydrophobic polypeptides. Therefore, although the structure maintains the path, the bias toward non-regulated protein growth in the hollow region suggests that the conventional bacteria-like lower part of the tunnel is unlikely to be functional.

To further examine the lower part of the tunnel, we prepared an accurate secondary structure diagram of the mt-rRNA based on the model coordinates (Supplementary Fig. 7). This was important because the sequence diverges substantially and therefore its alignment is not informative. The comparative analysis of the secondary structure diagrams showed that the 50-nucleotide long reduction of rRNA H24, which was reported to have ignited the remodeling of the tunnel path in the *S. cerevisiae* mitoribosome is also observed in our study (Fig. 5). This prompted us to analyze the region of the tunnel path that was shown to involve structural adaptations in *S. cerevisiae*. While the rRNA occupying this region is generally conserved, proteins uL22m and uL24m are extended in their N-termini so that two helical elements prevent accessibility (Fig 4c). Therefore, the yeast-like tunnel path is obstructed by the mitoribosomal proteins and is unlikely to be functional.

**Fig. 5:**
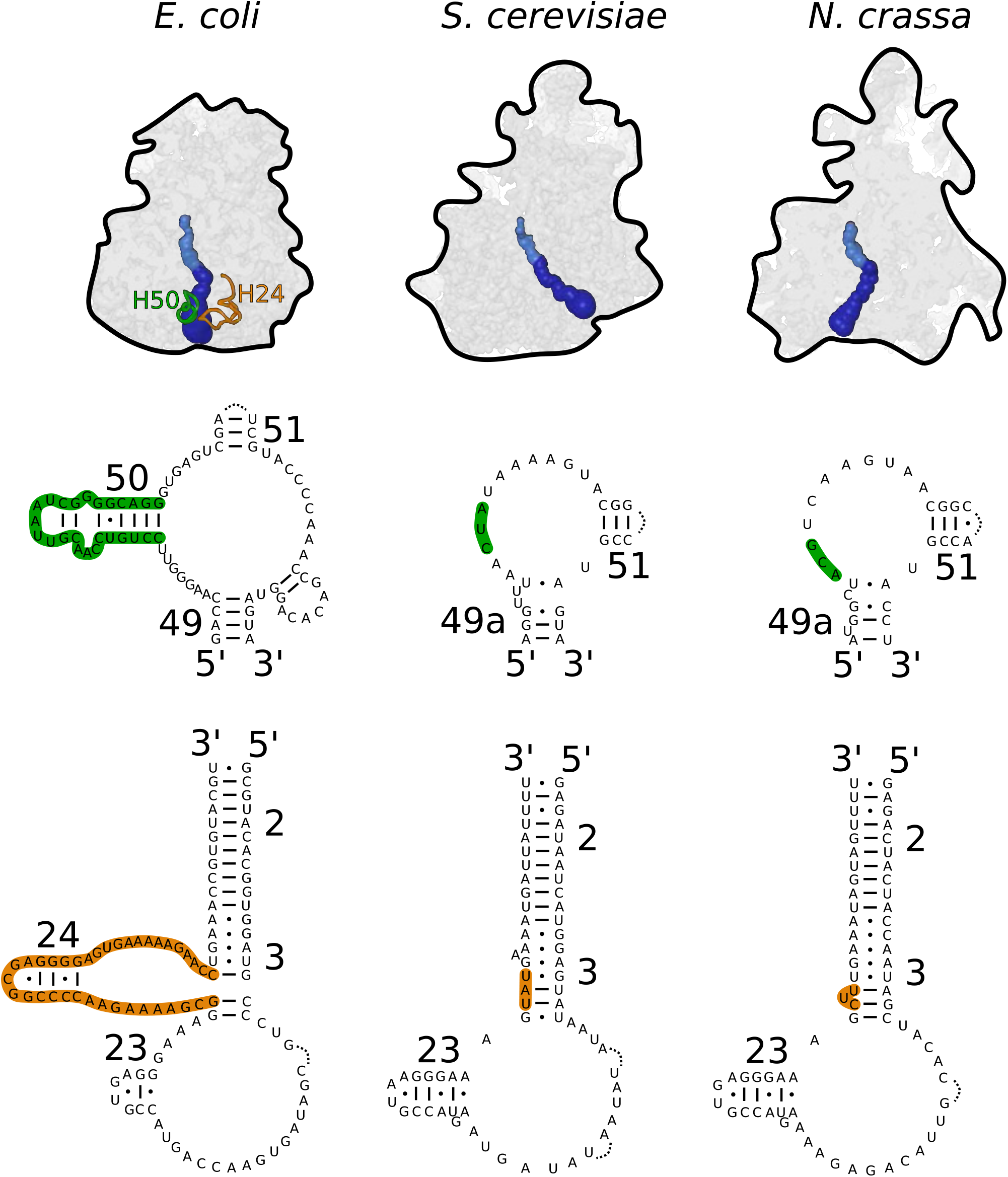
Comparison of the tunnel paths and the rRNA secondary structure elements involved in its formation. The different tunnel paths are shown for the bacterial ribosome (*E. coli*) and the mitoribosomes from *S. cerevisiae* and *N. crassa*. The locations of the rRNA helices H24 (orange) and H50 (green) are indicated. The secondary structure diagram for H24 and H50 regions features their deletion from the sequence of the mitochondrial DNA in *S. cerevisiae* and *N. crassa*.

Next, we inspected other remodeling of rRNA in the proximity to the tunnel. We identified a deletion of the 28-nucleotide long rRNA H50 segment ~58 Å from the start of the tunnel (Fig. 4d, Fig. 5). This remodeling gives rise to a channel that reaches the mitoribosomal surface ~30 Å away from the original tunnel. The 47-ES1 of the rRNA domain III and the proteins uL22m, uL23m, uL24m, and mL67 shape the boundaries of the tunnel exit leading to the mitoribosomal surface (Fig. 4d). Unlike the bacterial-like tunnel, it forms a wide path that would allow a higher degree of conformational sampling for a synthesized protein.

The exit site of this identified path is further accompanied with specific structural rearrangements. The rRNA element H59 was deleted, and H53 is unstructured and displaced, which is complemented by acquisition of the mitochondria-specific protein mL67/Mhr1. This mitoribosomal protein is also known as mitochondrial recombinase ^24^ that mediates mtDNA replication ^25^, repair ^26^, and was detected in the mitoribosomal assembly survey ^27^. It has a flexible extension of 25 residues, which is not present in *S. cerevisiae*, adjacent to the exit site and pointing towards the membrane plane (Supplementary Fig. 17c and Supplementary Fig. 18). Further, mitochondria-specific rRNA expansion segment 96-ES1 also points towards the membrane and the terminal 39 nucleotides appear to be disordered in our map. Together, the data suggest a potential adaptation of the tunnel exit (Supplementary Fig. 17c).

From the evolutionary perspective, the internal changes that have led to the deviation of the tunnel path are intrinsic to the rRNA fold. As in the case of the *S. cerevisiae* mitoribosome, the path was altered as a result of a deletion in rRNA. Since rRNA is encoded in the mitochondrial genome, it is the loss of genetic material from mitochondria that leads to a considerable effect on the mitoribosomal structure in the exit tunnel. This determines the geometry of the lower tunnel, which is reflected in the protein folding ^28^. Therefore, a further adaptation is required by the means of added protein (mL67/Mhr1) and rRNA (96-ES1) that gain function for the adjustment of the exit site in respect to the membrane. Consequently, our structure suggests that constructive neutral evolution of mitochondrial DNA ^29^ drives the architecture of the mitoribosomal exit tunnel.

### Visualization of mtLSU-Atp25 assembly intermediate suggests a regulatory mechanism

During 3D classification, we recognized a minor population of the particles corresponding to the isolated mtLSU, and subsequent processing resulted in 3.03 Å resolution reconstruction from 24,142 particles (Supplementary Fig. 5). Fitting the mtLSU model revealed an extra density that appears as a protein element forming a heterodimer with uL14m. In addition, rRNA at the interface (38-ES2, H63, H67-71, H80, 82-ES6, H101, PTC) appears as unfolded (Fig. 6). Further, the L1 stalk changes to a conformation that is more flexible and the density for bL9m next to L1 stalk is absent, whereas all the other mtLSU proteins are present (Fig. 6a).

**Fig. 6:**
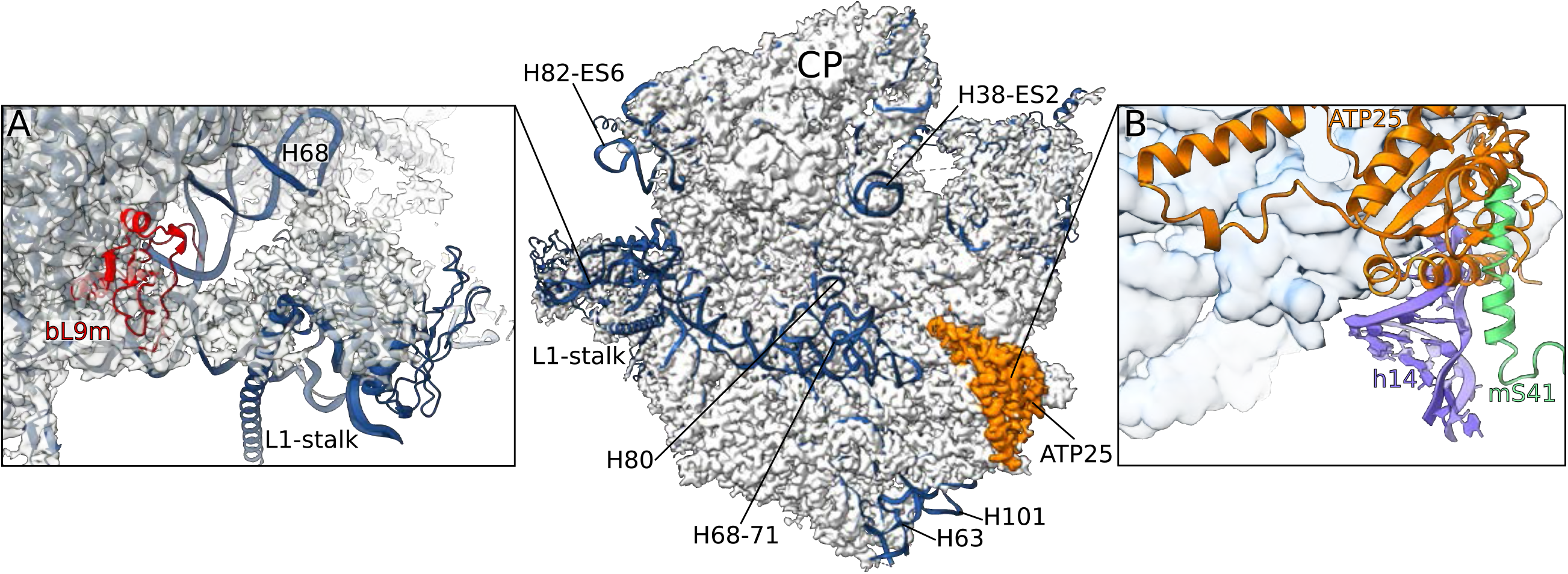
Structure and function of the mtLSU-Atp25 assembly complex. Superposition of the assembly intermediate cryo-EM map (gray) with the mature mtLSU model (blue) reveals unfolded interfacial rRNA. **a** conformational change of the L1 stalk, missing bL9m (red). **b** presence of Atp25 (orange) that would clash with the mtSSU h14 and mS41, preventing subunit association.

We identified the extra density as Atp25, an ortholog of the bacterial ribosome silencing factor (Rsf), for which there is no high-resolution structural information in complex with a ribosome ^30,31^. Its human mitochondrial counterpart, MALSU1, has been shown to involve late assembly stages ^12,32^. The gene *Atp25* codes for 699 amino acids, including the targeting sequence, an N-terminal part related to Rsf/MALSU1, and a C-terminal part called the M-domain. In *S. cerevisiae*, Atp25 is cleaved between the N- and C-terminal parts during mitochondrial import ^27^ and the M-domain forms a separate functional protein that stabilizes the mRNA of the ATP synthase subunit Atp9 ^33^. The gene fusion is suggested to have evolved in order to prevent off target effects of Atp25 on the cytosolic ribosomes prior to entry to the mitochondria ^27^. We observe only the Rsf/MALSU1-related part of Atp25 on the mitoribosome, confirming the function in the assembly of the mtLSU.

Atp25 presents only in the class with the missing bL9m and unfolded interfacial rRNA that affects the L1 stalk. In the mature mitoribosome, bL9m is anchored at the base of the L1 stalk near the E-site, which is stabilized through tertiary interactions with the rRNA. Therefore, the recruitment of bL9m requires specific L1 stalk conformation that is interdependent with rRNA folding, including the PTC.

Atp25 bound to uL14m sterically obstructs the binding of the mtSSU by preventing the formation of bridge B8 with its globular domain (Fig. 6b). The steric block spans ~25 Å and would clash with h14 of rRNA and the N-terminus of mS26 of the mtSSU. Therefore, the eviction of Atp25 must take place during the final stages of maturation to alleviate the steric hindrance on subunit joining. A similar mechanism was proposed for eIF6 in the cytoplasmic translation apparatus, where protein factor SBDS senses the structural integrity of the functional sites before the displacement ^34^. Our data imply that since this is a prerequisite step for the folding of interfacial mt-rRNA into a native-like conformation, it is also a requirement for the binding of bL9m. This shows that bL9m attachment is mechanistically regulated during the assembly of the mitoribosome.

## Conclusions

This study reveals a dedicated mitoribosomal platform for mRNA loading and a protein mL108 that is associated with the L1 stalk and involved in tRNA translocation. The distinct exit tunnel suggests that the evolution of the nascent chain emergence in mitochondria is driven primarily by changes in the rRNA that is encoded in the mitochondrial genome in all species. Finally, the described assembly intermediate lacking bL9m proposes a regulatory mechanism.

## Methods

### Purification of mitoribosomes

Mitochondria were isolated from *Neurospora crassa* strain K5-15-23-1 ^36^ which contains a His-tagged version of Oxa1. The *N. crassa* mycelia were grown in aerobic conditions using 10 L of the Vogel’s growth medium ^37^, supplemented with L-lysine and L-leucine at 25°C for 16 h. Each growth flask was supplemented with 0.1% chloramphenicol 1 h prior to harvesting. To separate the moisture from the mycelia, the culture was filtered through muslin. The dry mycelia were transferred to a pre-cooled mortar and pestle to lyse the cells. All further operations were performed at 4°C. The cells were lysed using sea-sand (silicon dioxide) and SEMP buffer (250 mM sucrose, 1 mM ethylenediaminetetraacetic acid (EDTA), 10 mM MOPS-KOH pH 7.2, 2 mM phenylmethanesulfonyl fluoride (PMSF), and 0.2% chloramphenicol), by applying a 20 min grinding course. Sand and cell debris were removed using differential centrifugation at low-speed (2000 x g). A subsequent high-speed centrifugation step (17,500 x g) was carried out to sediment crude mitochondria. The mitochondrial pellet was then resuspended in SEM buffer (250 mM sucrose, 1 mM EDTA, and 10 mM MOPS-KOH pH 7.2). To further purify, the crude mitochondria were undergone a sucrose gradient in SEM buffer for 1h at 139,400 x g with an SW40 Ti rotor (Beckman Coulter). Mitochondrial band was pooled and stored at −80 °C.

Mitochondria were lysed in 4 volumes of lysis buffer (25 mM Hepes-KOH pH 7.5, 20 mM KCl, 15 mM Mg(OAc)_2_, 2% *n*-dodecyl-β-D-maltoside (DDM), 0.0075 % cardiolipin, 2 mM dithiothreitol (DTT), 1 tablet of protease inhibitor (cOmplete™, Roche), and RNase inhibitor (RNaseOUT, Invitrogen)) and incubated for 5 min at 4°C. The membrane was separated by centrifugation of the mitochondrial lysate at 30,000 x g for 20 min followed by a second centrifugation step with the same settings. The supernatant was loaded onto a 1.0 M sucrose cushion in the resuspension buffer (20 mM Hepes-KOH pH 7.5, 15 mM KCL, 15 mM Mg(OAc)_2_, 0.05% DDM, 0.0075% cardiolipin, and 2 mM DTT) with RNase inhibitor and centrifuged at 237,100 x g with a 70 Ti rotor for 4 h. The pellet was then resuspended in the resuspension buffer, loaded on a 15-30% sucrose gradient in the resuspension buffer, and centrifuged at 94,100 x g with an SW40 Ti for 16 h at 4°C. Fractions containing mitoribosomes were collected and the mitoribosomes were pelleted by centrifugation at 356,000 x g with a TLA120.2 rotor (Beckman Coulter) for 45 min at 4°C and resuspended in the resuspension buffer.

### Cryo-EM and image processing

Freshly purified mitorobosome sample (A_260_ = 3.0) was used for grid preparation. 3 μl aliquots of purified mitoribsomes was incubated for 30 s on glow-discharged holey carbon grids (Quantifoil R2/2, 300 mesh, copper) pre-coated with a home-made continuous carbon film with thickness of ~ 27 Å. The grids were thereafter vitrified in a liquid ethane with a Vitrobot MKIV (FEI/Thermo Fisher) using a blotting time of 3 s at 4°C and 100% humidity. Micrographs were collected with a 300 kV Titan Krios (FEI/Thermo Fisher) transmission electron microscope, 70 μm C2 aperture, using a slit width of 20 eV on a GIF-Quantum energy filter (Gatan). A K2 summit detector (GATAN) was used in the counting mode at the calibrated magnification of 130,000 (yielding a pixel size of 1.06 Å). An exposure time of 9 s yielding a total of 35 e/Å^2^ was dose fractionated into 20 frames. In total 3,172 movies were collected automatically during two consecutive days using EPU (FEI/Thermo Fisher) data collection software. Defocus values ranged between 0.8 and 3.0 μm.

Collected movie frames were aligned and corrected for both global and non-uniform local beam-induced movements using MotionCor ^38^ and the contrast transfer function (CTF) parameters were estimated using Gctf ^39^, inside the SCIPION program ^40^. Subsequent data processing steps were performed using RELION-2.1 and 3.0 ^41^. First, 647 particles were manually picked, followed by two-dimensional (2D) classification. Four good 2D class averages were used for reference-based picking for the second round. 265,710 picked particles were subjected to 2D classification and 50 good 2D classes were selected (Supplementary Fig. 1). Retained 223,605 particles were classified into six classes by three-dimensional (3D) classification, resulted in four good mito-monosome classes (131,806 particles), one class with weak mtSSU density (36,053 particles), and one low quality class containing poorly-aligned or broken particles. From the class with weak mtSSU density, isolated mtLSU particles (24,142 particles) are classified out by further 3D classification. Pooled good particles were subjected to 3D auto-refinement. Per-particle CTF refinement ^42^, followed by Bayesian polishing ^43^ and the second round of per particle CTF refinement to improve resolution, resulted in reconstructions of mito-monosome and the isolated mtLSU with 2.83 and 3.03 Å resolution, respectively Supplementary Fig. 1). Resolution is estimated based on the gold standard criterion of Fourier shell correlation (FSC) = 0.143 between the reconstructed two half maps.

Due to the relative movement between the mtSSU and mtLSU in monosome, we suffered from low resolution in the mtSSU. We therefore decided to improve the quality of the maps by focused masked refinement for the mtSSU and mtLSU separately. We obtained the masked maps of mtLSU and mtSSU with 2.74 and 2.85 Å resolution, respectively. Further, improvement of local resolutions was achieved by focused refinement using the local masks as described in the Supplementary Material.

All the density maps were locally resolution filtered by applying a negative B-factor estimated automatically by RELION-3.0.

The observed motion between mtLSU and mtSSU prompted us to separate possible sub-states of the monosome. Firstly, we classified the particles in two major classes (rotated and non-rotated states) by overall 3D classification. To facilitate classifying tRNA states, signal subtraction of ribosome was performed using a cylindrical mask covering the A-, P-, and E-sites for the rotated and non-rotated states separately by RELION 3.0, following by focused 3D classification without alignment. For the rotated state, the same cylindrical mask was applied for the classification, which separated the P/E hybrid tRNA bound mitoribosomes (37,908 particles). For non-rotated state, smaller masks were needed to classify tRNAs. The P/P tRNA bound mitoribosomes were separated applying a mask covering only the P-site, while E/E tRNA bound ones were separated by a mask covering only the E-site. The P/P tRNA and E/E tRNA bound mitoribosomes were 24,611 and 23,802 particles, respectively. Among them, only 4,136 particles are overlapping and have both P/P and E/E tRNAs, which are too few for high-resolution reconstruction. Local resolution of the tRNA bound mitoribosomes were also improved by using local masked refinement as described in the Supplementary Material.

### Model building and refinement

Model building was carried out in Coot ^44^. Rigid body docking was carried out in UCSF Chimera ^45^. The model of the *S. cerevisiae* mitoribosome (PDB ID: <u>5MRC</u>) was used as a reference. For proteins whose orthologs existed in *S. cerevisiae* structure, the homology models of *N. crassa* were generated by SWISS Model ^46^, placed into the density, manually revised and built unmodeled and unique regions, and real space fit in *Coot*. Previously unknown or unmodeled proteins were modeled *de novo* in *Coot*. For the correct assignment of the proteins, sequence stretches of reasonable quality were directly identified from the map and then applied to BLAST ^47^ and/or compared to the protein sequences gained from mass spectrometry (Supplementary Data 1). The ribosomal protein mL108 was named according to the standard nomenclature widely accepted in the ribosome community ^48^. The number was chosen in a chronological order of discover also considering unpublished data. The entire rRNAs, tRNAs, and mRNAs were modeled *de novo* in *Coot*. Since bound tRNAs and mRNAs are mixture of any mitochondrial ones, each residue was assigned as either A, U, G, or C, based on the density and conservation. Ligands and metal ions were placed into the density.

For model refinement of the consensus mtLSU and mtSSU and the monosomes in the three tRNA states, the local masked refined maps with B-factor sharpening and local-resolution filtering were merged using the program phenix_combine_focused_maps in the Phenix software suite ^49^, thereby generating a map for model refinement. For the isolated mtLSU, the overall refined map with B-factor sharpening and local-resolution filtering was used for model refinement. Refinement of all models was performed using Phenix.real_space_refine ^50^. The final structure was validated using the MolProbity ^51^ implementation of the Phenix suite.

### Figure preparation

All structural figures were generated with PyMOL ^52^, UCSF Chimera ^45^, and ChimeraX ^53^. The secondary structure diagrams of the 16S and 23S rRNA were generated using XRNA (http://rna.ucsc.edu/rnacenter/xrna/xrna.html). The shape of the polypeptide tunnels were calculated with the software Mole^54^.

## Acknowledgements

We thank Frank Nargang for providing the *N. crassa* strain, and Marta Carroni and Julian Conrad for supporting cryo-EM data collection. This work was supported by the Swedish Foundation for Strategic Research (FFL15:0325), Ragnar Söderberg Foundation (M44/16), Swedish Research Council (NT_2015-04107), Cancerfonden (2017/1041), European Research Council (ERC-2018-StG-805230), Knut and Alice Wallenberg Foundation (2018.0080), EMBO Young Investigator Program. Y.I. is supported by H2020-MSCA-IF-2017 (799399-Itohribo). The data was collected at the SciLifeLab cryo-EM facility funded by the Knut and Alice Wallenberg, Family Erling Persson, and Kempe foundations. Mass spectrometry analysis was performed by the clinical proteomics mass spectrometry facility of the Karolinska University Hospital and SciLifeLab proteogenomics core facility. We thank John Walker and Alan Brown for constructive comments, and Katrina Albert for proofreading the manuscript.

## Author contributions

A.A. designed the project. N.M prepared the sample and collected cryo-EM data. Y.I., A.N. and N.M. processed the data and built the model. Y.I., A.N. and A.A. wrote the manuscript with help from N.M. and J.M.H.

## Data availability

Cryo-EM maps have been deposited at the EMDB with the following accession codes: <u>EMD-10958</u>, <u>EMD-10961</u>, <u>EMD-10962</u>, <u>EMD-10963</u>, <u>EMD-10965</u>, <u>EMD-10966</u>, <u>EMD-10967</u>, <u>EMD-10968</u>, <u>EMD-10969</u>, <u>EMD-10970</u>, <u>EMD-10971</u>, <u>EMD-10972</u>, <u>EMD-10973</u>, <u>EMD-10974</u>, <u>EMD-10975</u>, <u>EMD-10976</u>, <u>EMD-10977</u>, <u>EMD-10978</u>, <u>EMD-10979</u>, <u>EMD-10980</u>, <u>EMD-10981</u>, <u>EMD-10982</u>, <u>EMD-10983</u>, <u>EMD-10984</u>, <u>EMD-10985</u>, <u>EMD-10986</u>, <u>EMD-10988</u>, <u>EMD-10989</u>, <u>EMD-10990</u>, <u>EMD-10991</u> and <u>EMD-10992</u>. The atomic coordinates have been deposited at the PDB with the following accession codes: <u>6YWS</u>, <u>6YW5</u>, <u>6YWX</u>, <u>6YWY</u>, <u>6YWE</u> and <u>6YWV</u>.

## Supplementary Information

**Supplementary Fig. 1:**
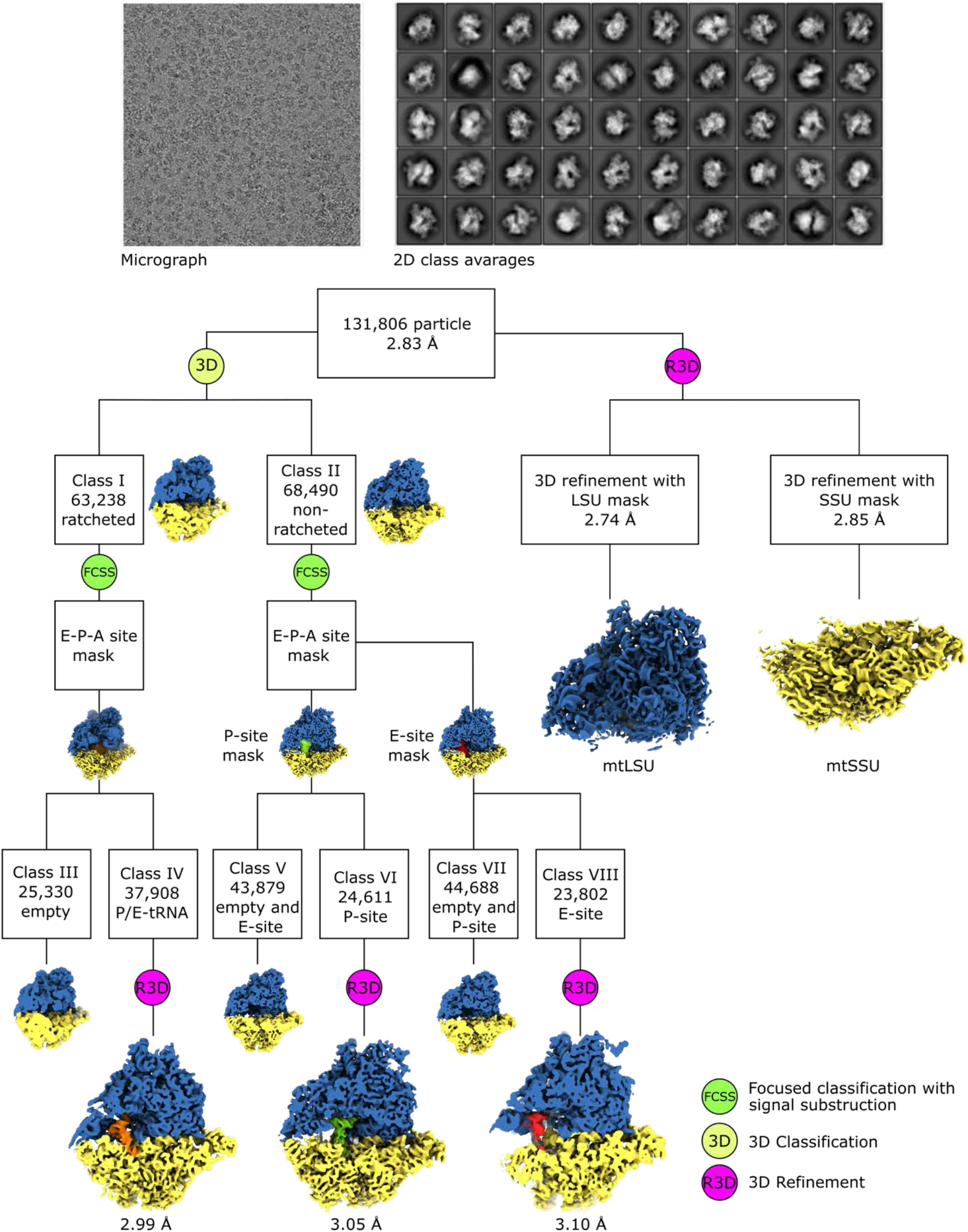
Cryo-EM data collection and processing scheme. Representative micrograph and 2D class averages. Data refinement protocol and classification scheme for mtLSU, mtSSU and monosome in three tRNA bound states.

**Supplementary Fig. 2:**
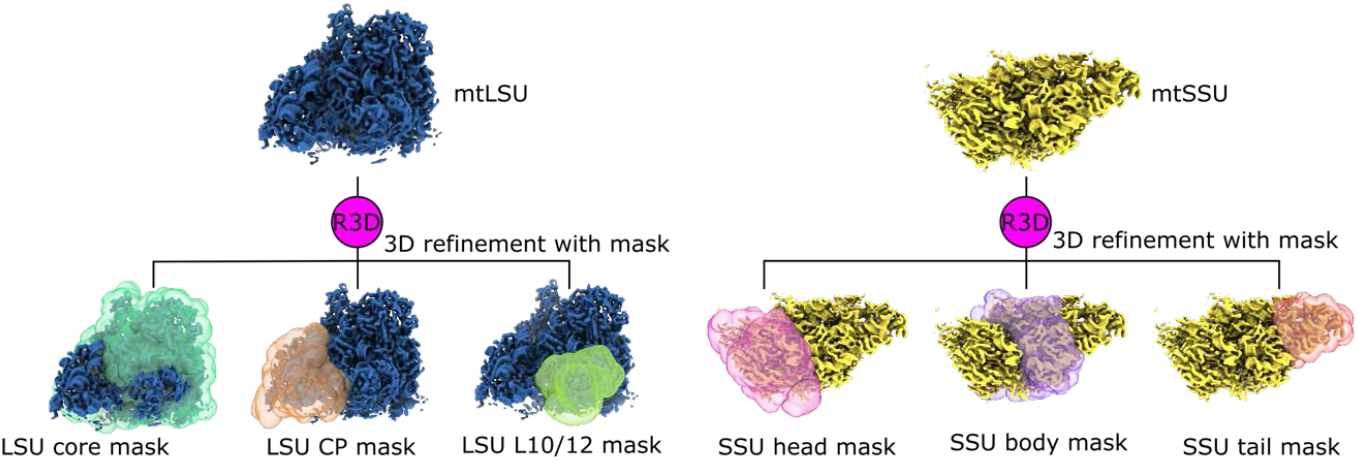
Masked refinement of mitoribosomal subunits. Masks for mtLSU core, CP, L7/12 stalk; mtSSU head, body, tail were used to improve the density locally.

**Supplementary Fig. 3:**
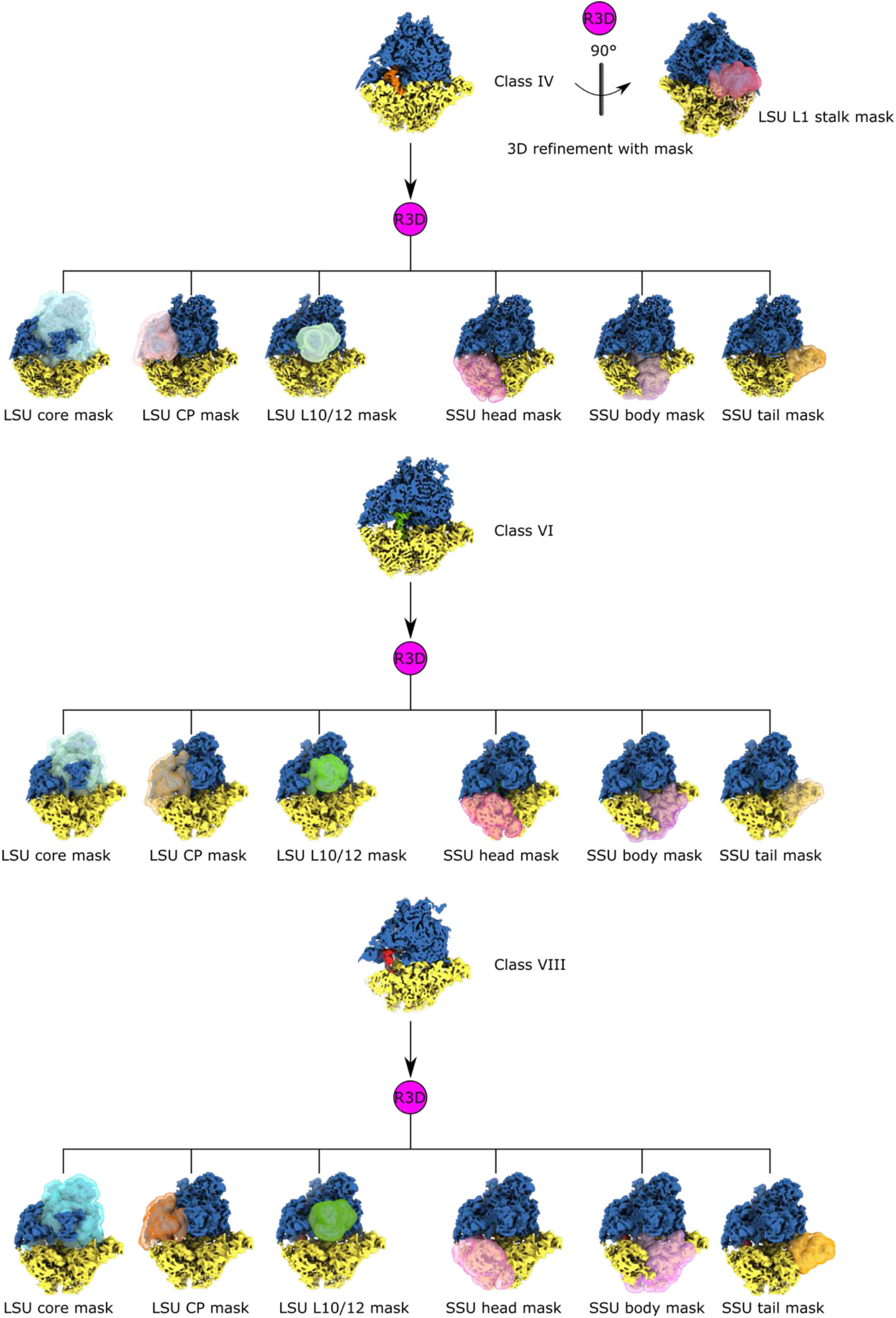
Masking refinement of the mitoribosomes in tRNA bound states.

**Supplementary Fig. 4:**
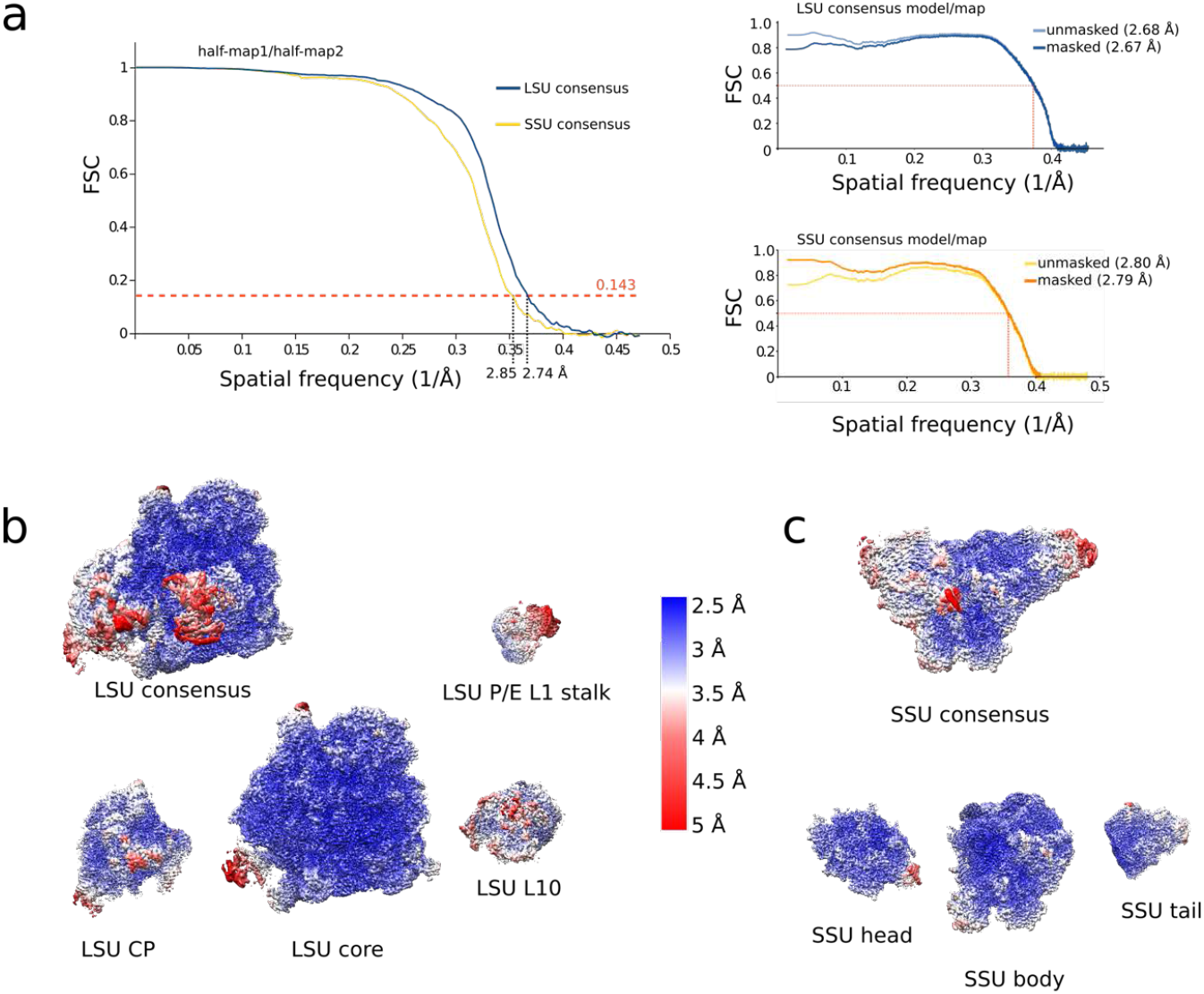
Overall and local resolution estimations of mitoribosomal subunits. **a** Fourier Shell Correlation (FSC) curves of half-maps and map-to-model are shown. Cut-off values of 0.143 (gold-standard) and 0.5 (model-to-map) are indicated and the corresponding resolution estimates are shown **b** mtLSU cryo-EM maps viewed according to local resolution. **c** mtSSU cryo-EM maps viewed according to local resolution.

**Supplementary Fig. 5:**
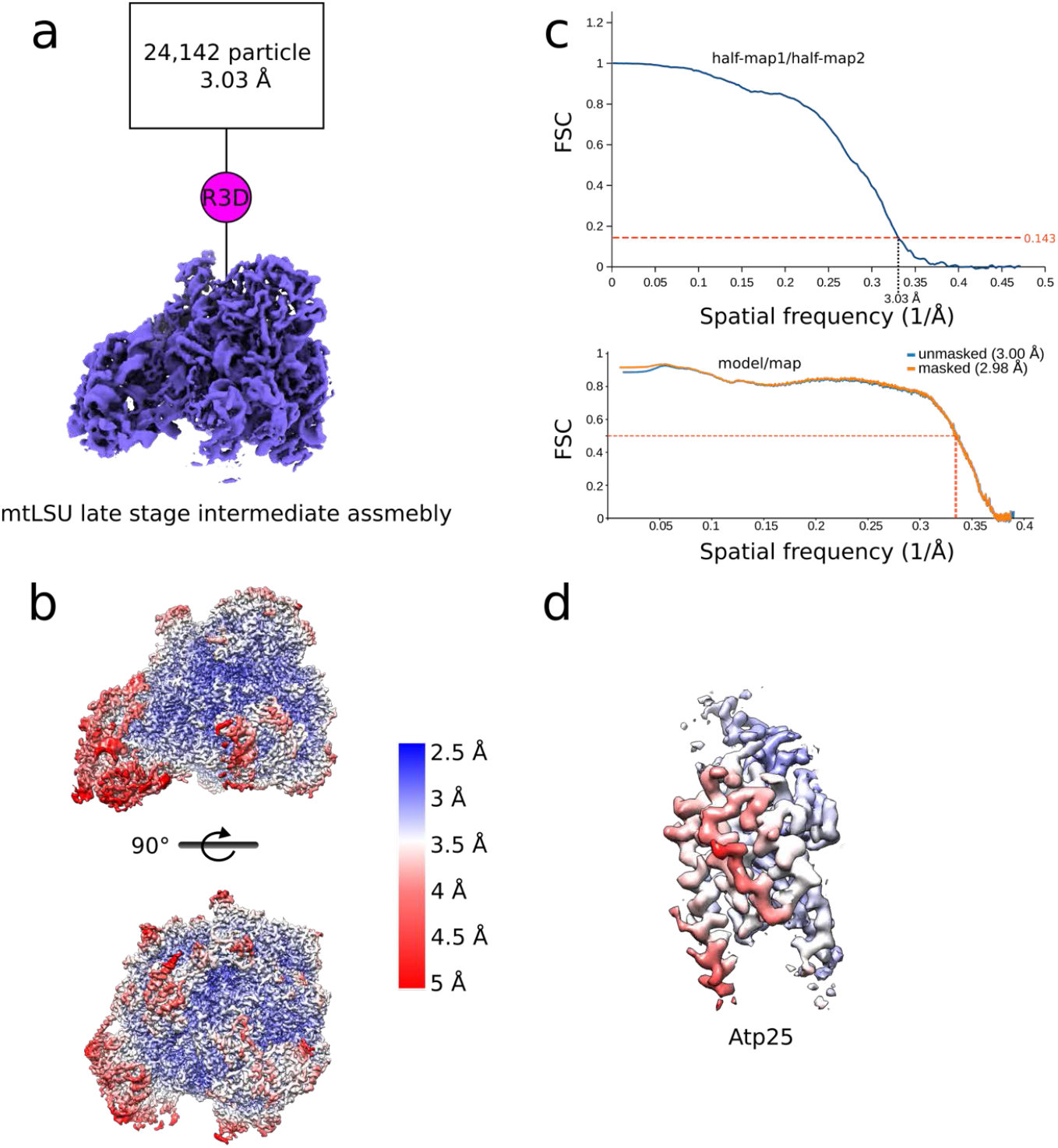
Data processing and overall and local resolution estimations of mtLSU-Atp25 assembly intermediate. **a** Processing scheme. **b** Fourier Shell Correlation (FSC) curves for half-maps and map-to-model are shown. Cut-off values are indicated and resolution estimates are shown. **c** mtLSU-Atp25 assembly intermediate cryo-EM maps colored according to local resolution. **d** Zoomed in map for Atp25 colored by local resolution.

**Supplementary Fig. 6.**
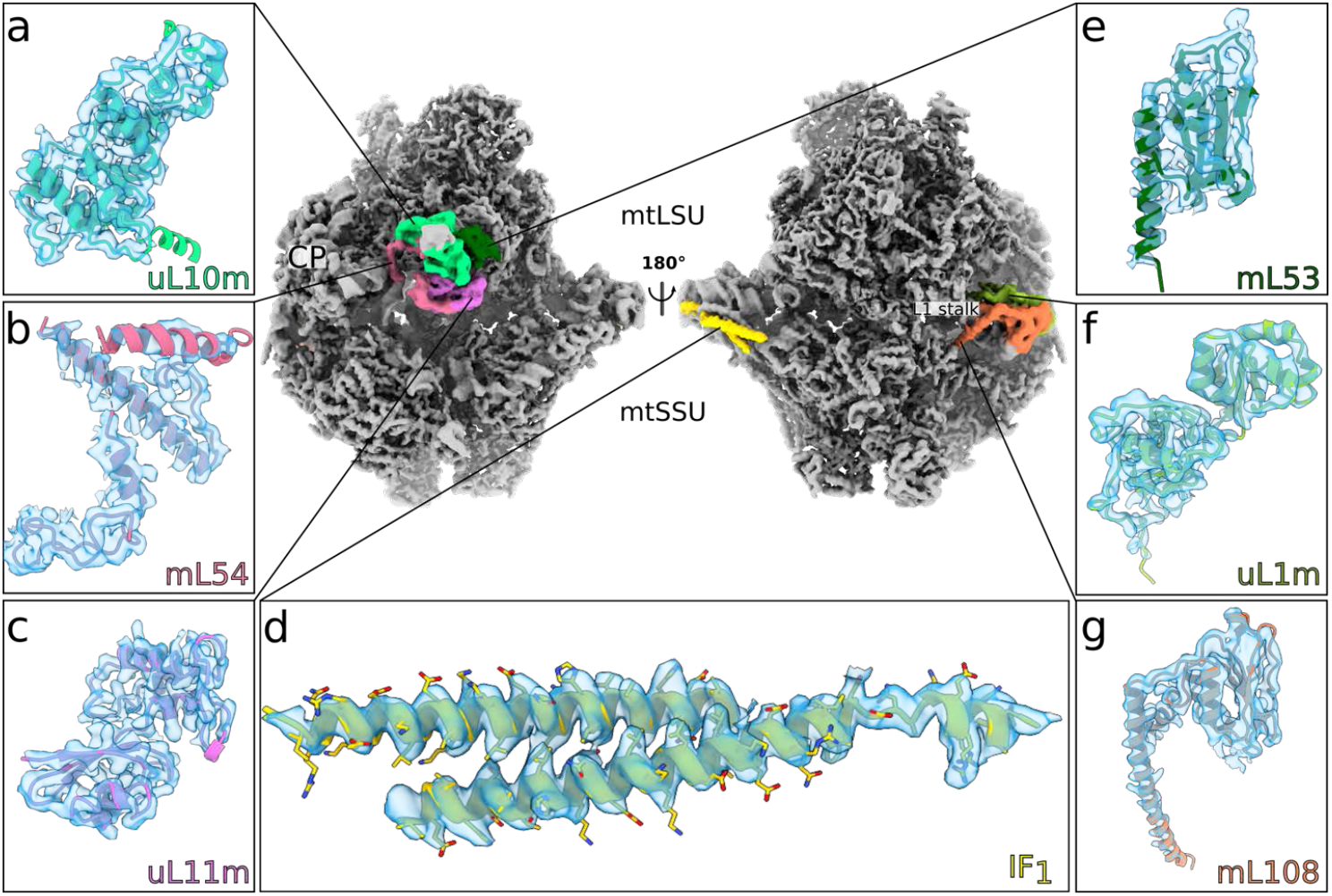
Cryo-EM reconstruction and newly modelled proteins. Two views of a composite cryo-EM map with newly identified proteins colored. The close-up views show the newly modeled proteins with their corresponding density map: **a** uL10m. **b** mL54. **c** uL11m. **d** IF_1_. **e** mL53. **f** uL1m. **g** mL108.

**Supplementary Fig. 7:**
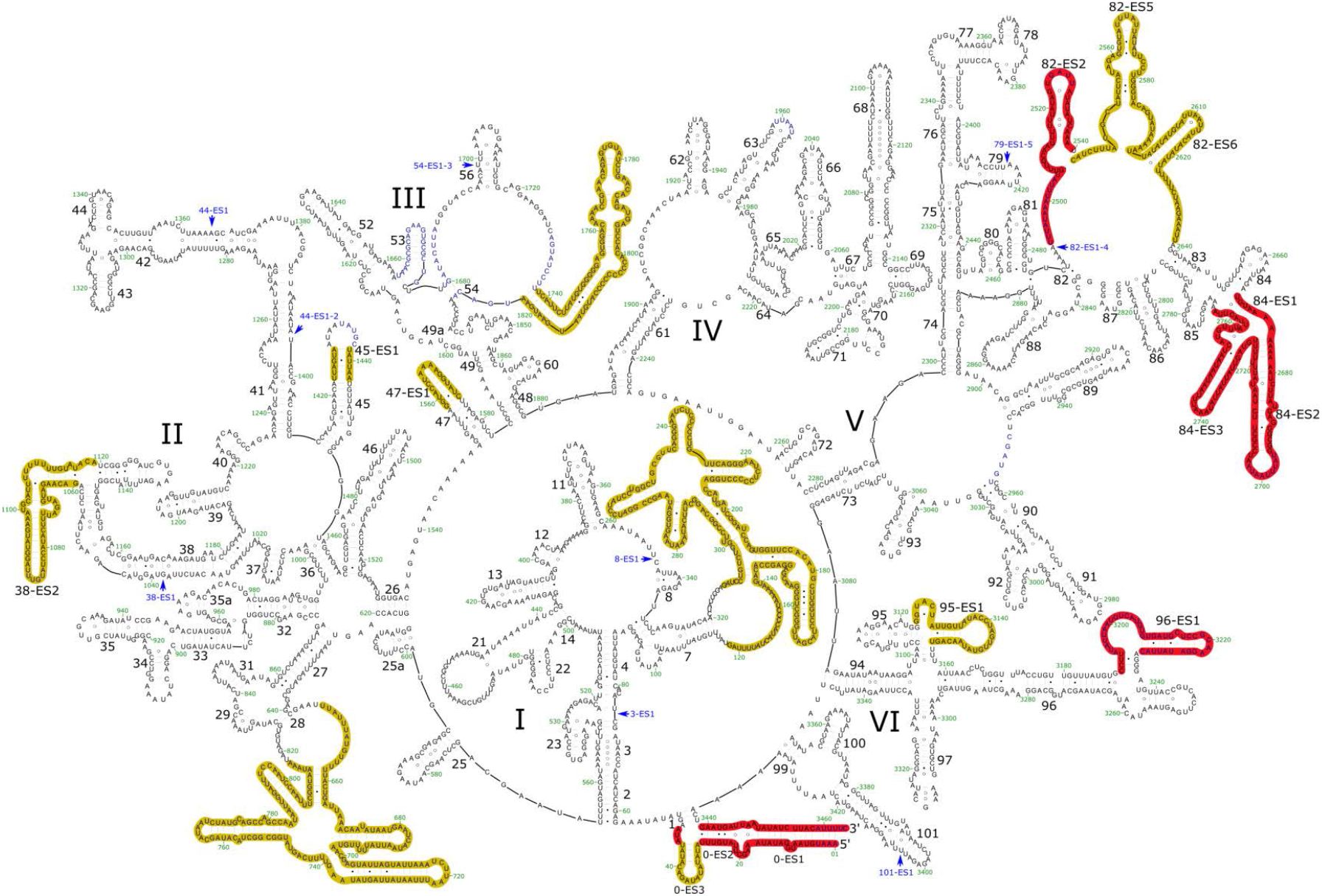
Structure-based secondary structure diagram of the mtLSU rRNA. Expansion segments shared between *N. crassa* and *S. cerevisiae* are highlighted with red, specific expansion segments are highlighted with yellow. Those present in *S. cerevisiae* only are indicated by light blue arrows. rRNA domains are labeled with roman numerals.

**Supplementary Fig. 8:**
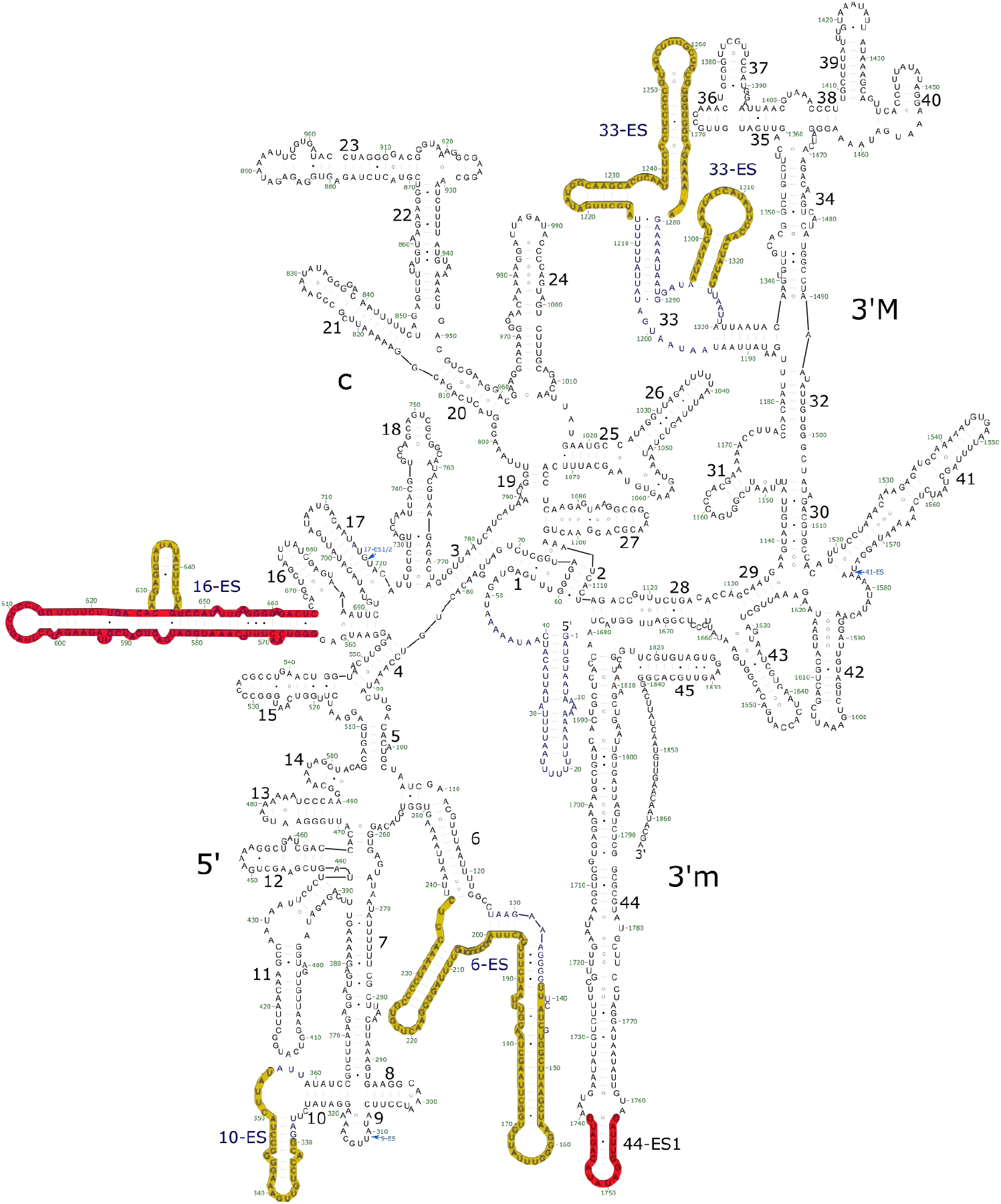
Structure-based secondary structure diagram of the mtLSU rRNA. Expansion segments shared between *N. crassa* and *S. cerevisiae* are highlighted with red, specific expansion segments are highlighted with yellow. Those present in *S. cerevisiae* only are indicated by light blue arrows. rRNA domains are labeled with roman numerals.

**Supplementary Fig. 9:**
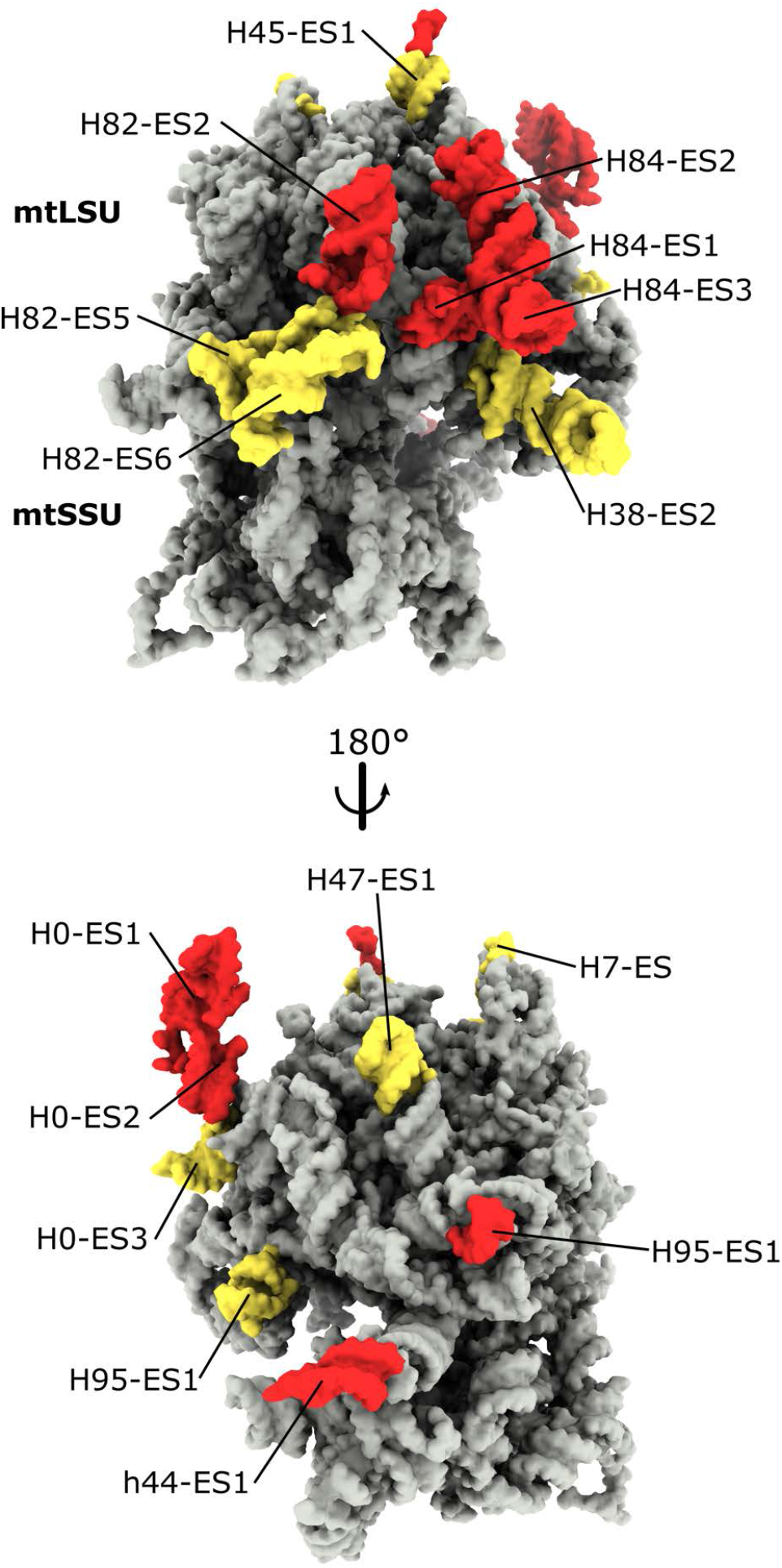
3D view of the rRNA of mtLSU and mtSSU. The conserved rRNA is in gray. ESs shared with *S. cerevisiae* are in red, and specific ESs in yellow.

**Supplementary Fig. 10:**
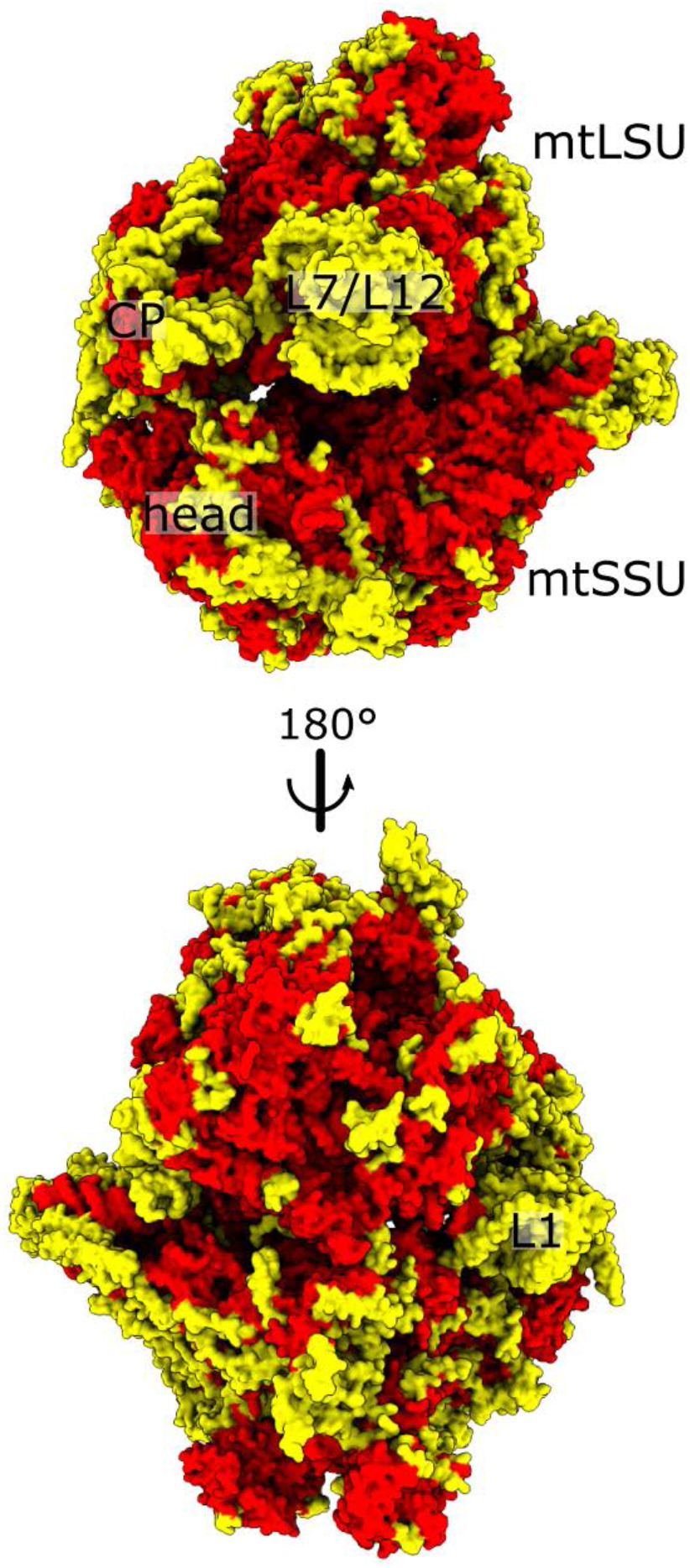
Improvements in the fungal mitoribosomal model. The newly modeled and improved features in the current structure (yellow) compared with the previous model (red) of the *S. cerevisiae* mitoribosome (PDB ID: <u>5MRC</u>).

**Supplementary Fig. 11:**
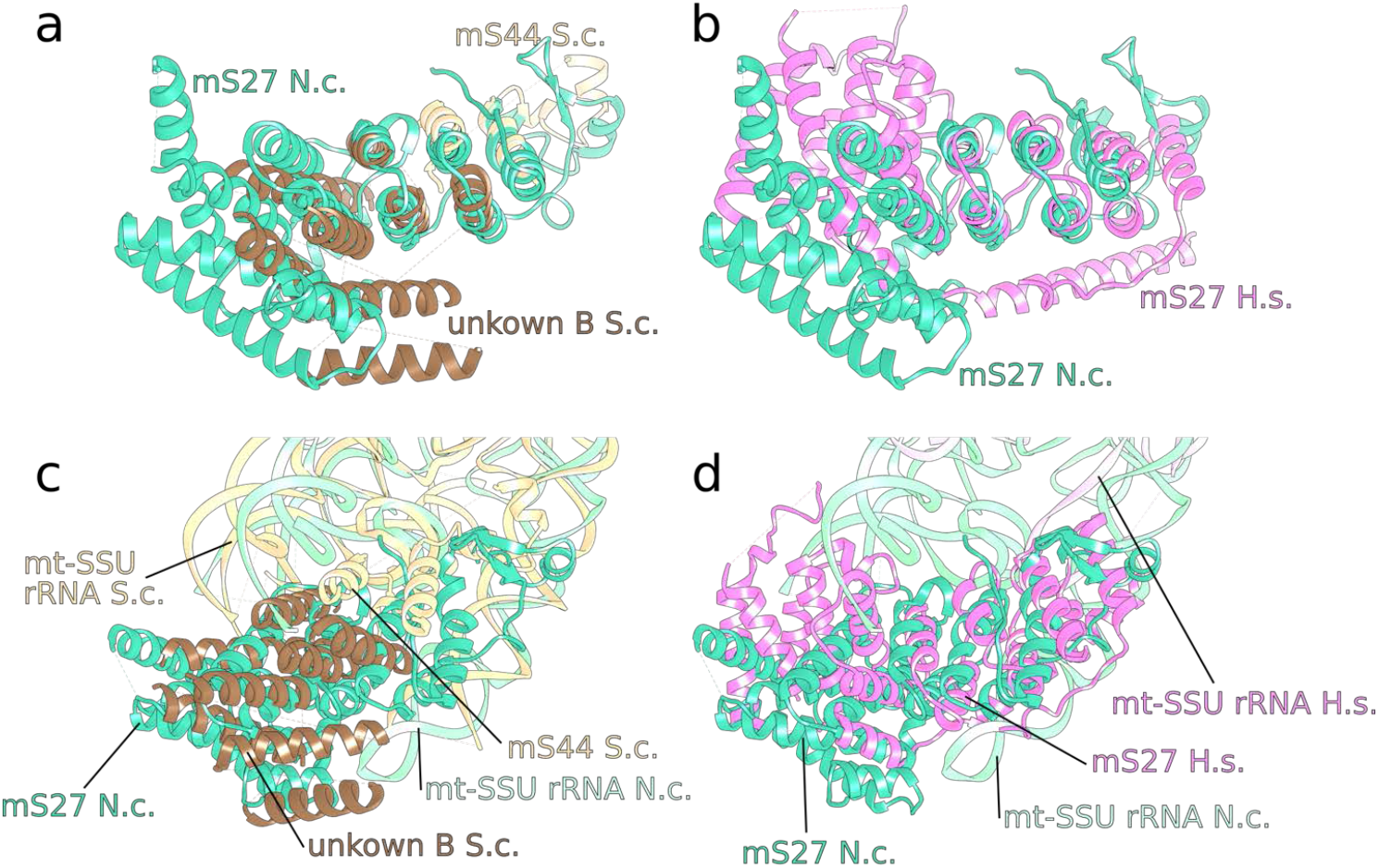
Superposition of mS27 orthologs. **a** *N. crassa* and *S. cerevisiae* mS27 is superimposed (previously assigned as mS44, ‘unknown B’). **b** *N. crassa* and *H. sapiens* mS27 is superimposed, showing similar fold. **c** Superposition based on mtSSU rRNA suggests the same location form mS27 from *N. crassa* and *S. cerevisiae*. **d** Superposition based on mtSSU rRNA suggests the same location form mS27 from *N. crassa* and *H. sapiens*.

**Supplementary Fig. 12:**
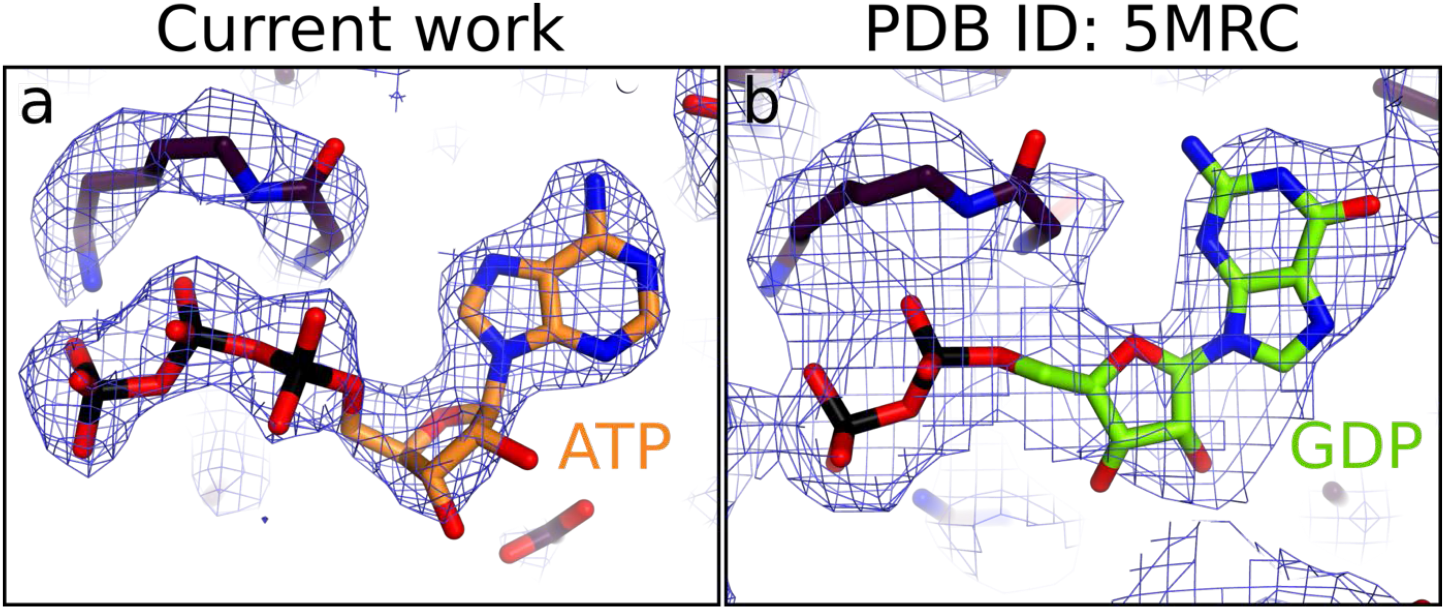
Comparison of the nucleotide density for mS29 between the current model of *N. crassa* and previous work. **a** The density for ATP bound to mS29 in the current work with the corresponding model **b** For comparison the density of the previously modeled GDP in *S. cerevisiae* is shown.

**Supplementary Fig. 13:**
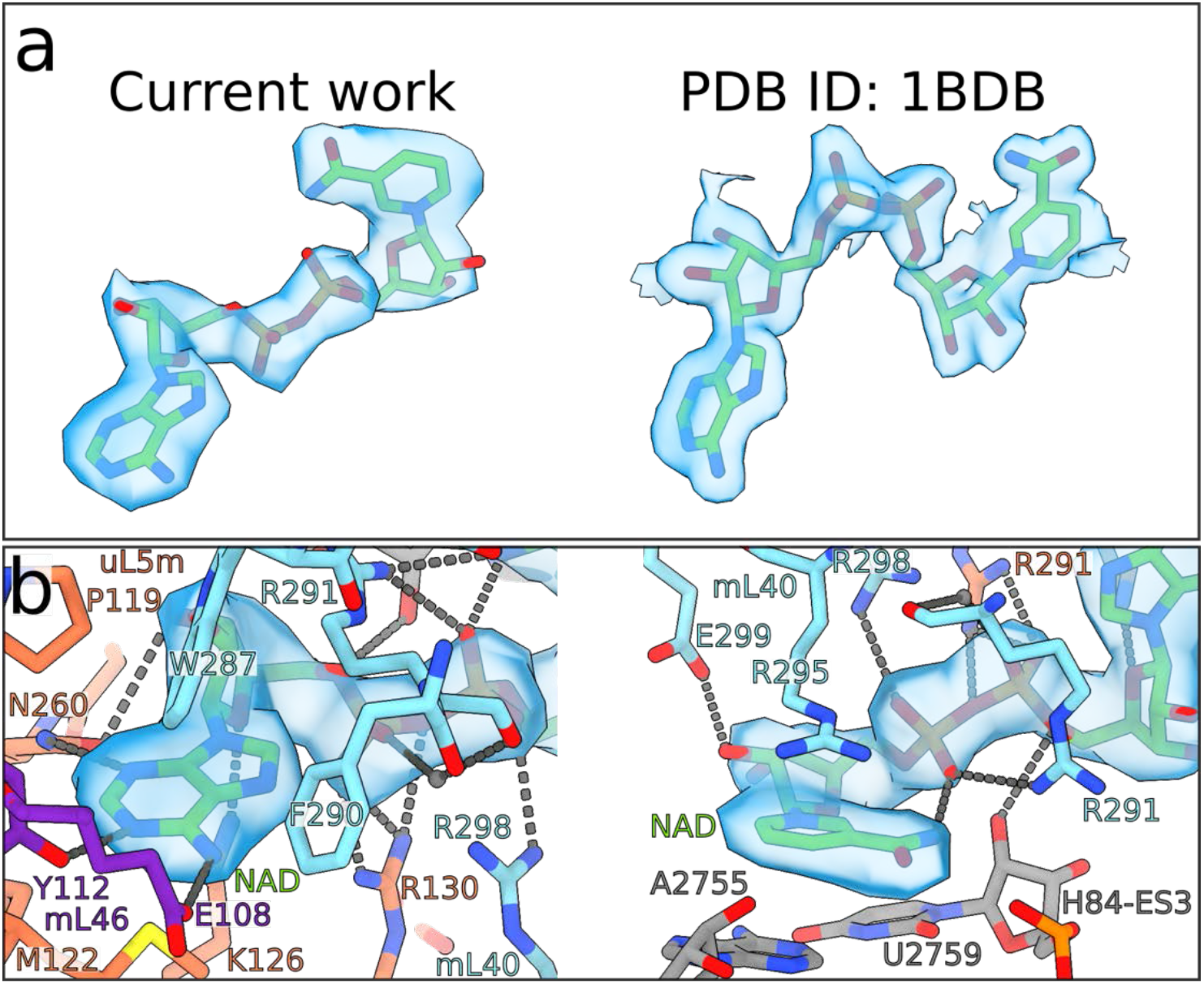
Density map and interactions of the NAD binding pocket in the CP of *N. crassa*. **a** Comparison of the density in the current work with 2 Å resolution crystal structure (PDB ID: <u>1BDB</u>) confirms the correct assignment of NAD in the model. Both studies show the nicotine amid in syn-conformation. **b** Interactions formed by NAD with uL5, mL40 and H84-ES3 rRNA showed from two opposite views. One view is centered on the adenine ring and the other view around the nicotine amid. Hydrogen bonds and charged interactions are indicated by black dashed lines. Residues and nucleotides are colored and labeled correspondingly.

**Supplementary Fig. 14:**
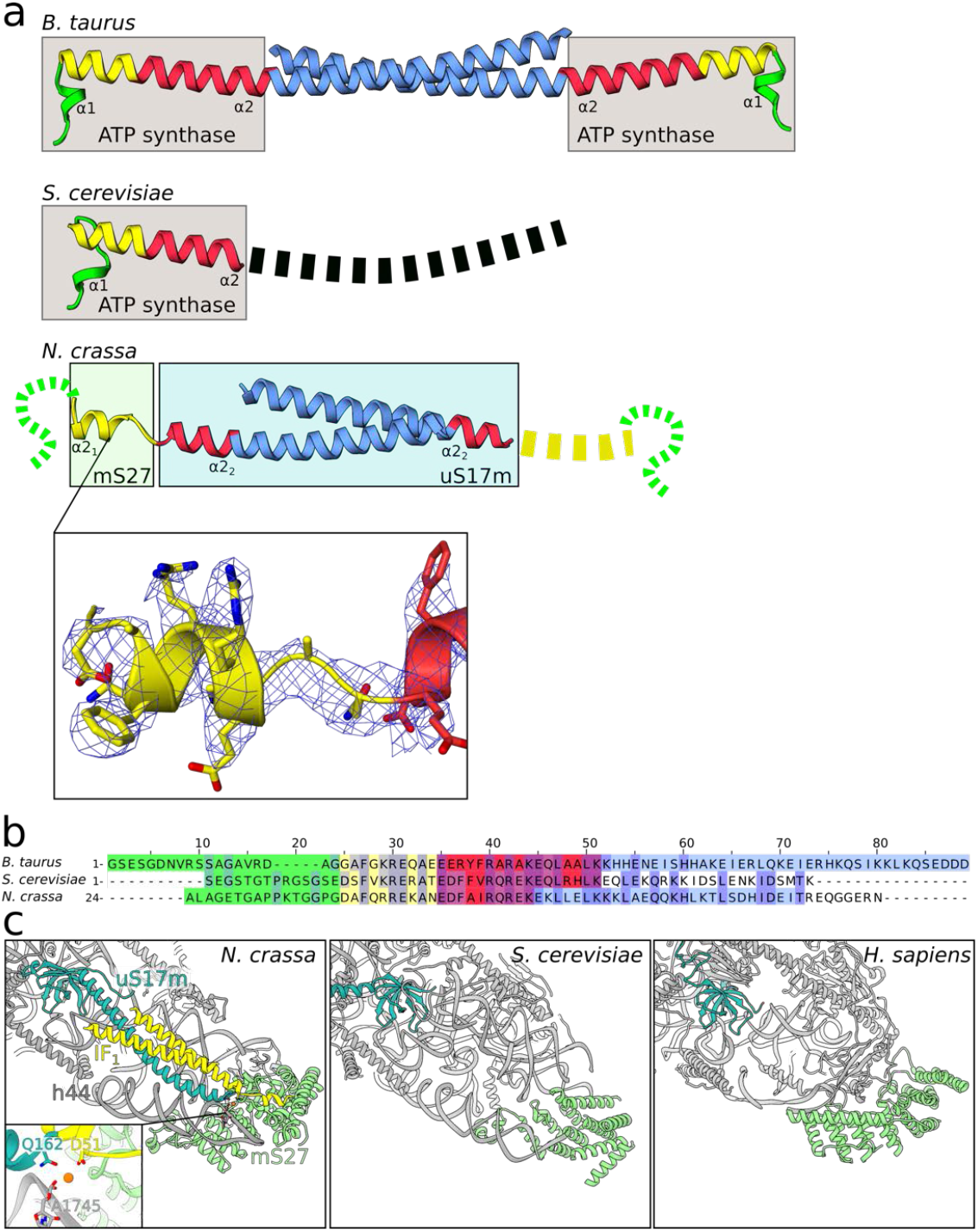
IF_1_ topology, structure, binding and conservation. **a** Comparison with *Bos taurus* (generated from PDB IDs: <u>1GMJ</u> ^1^ and <u>4TSF</u> ^2^) and *S. cerevisiae* (PDB ID: <u>3ZIA</u>). Density map with model is shown to illustrate the quality of the data in this region. The mitoribosome and F_1_-ATPase binding parts are indicated with boxes. For *N. crassa*, the disordered N-terminal region (residues 24-39), the short helix α2_1_ with a loop (residues 40-49), the longer helix α2_2_ outside the dimerization part (residues 50-58), and α2_2_ within the dimerization part (residues 59-88) are green, yellow, red, and blue, respectively. Corresponding residues are colored accordingly in *B. taurus* and *S. cerevisiae*, where α1 is located in the N-terminal region and α2_1_ and α2_2_ are a continuous helix α2. **b** Structure-based sequence alignment, colored as in **a**. The mitochondria targeting sequence is included in the residue numbering for *N. crassa*. **c** Structure of the IF_1_-binding site of *N. crassa* mtSSU and the corresponding regions from *S. cerevisiae* and *H. sapiens*. IF_1_, uS17m, and mS27 are indicated. The IF_1_ homodimer forms a helical bundle with the specific C-terminal extension of uS17m. The Mg-ion mediated bridge is shown in a zoom-in window.

**Supplementary Fig. 15:**
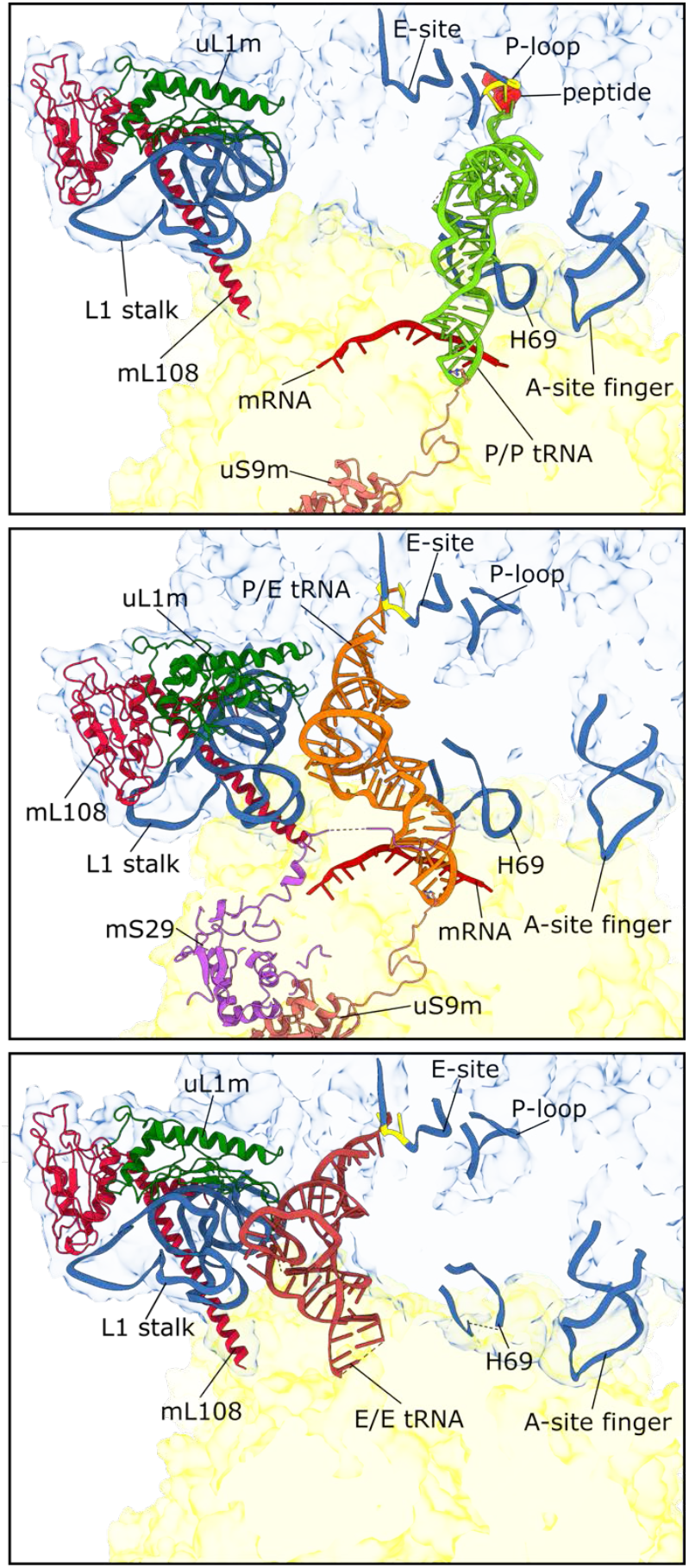
The L1-stalk and translocation of tRNA. The three states are structurally aligned on mtLSU. Transparent surfaces of mtLSU and mtSSU are colored in blue and yellow respectively. Cartoon representations of involved parts are shown and labeled. The P/P-tRNA state with the tRNA colored in green (top). The P/E-tRNA state with the tRNA colored in orange (middle). The E/E-tRNA state with the tRNA colored in red (bottom).

**Supplementary Fig 16:**
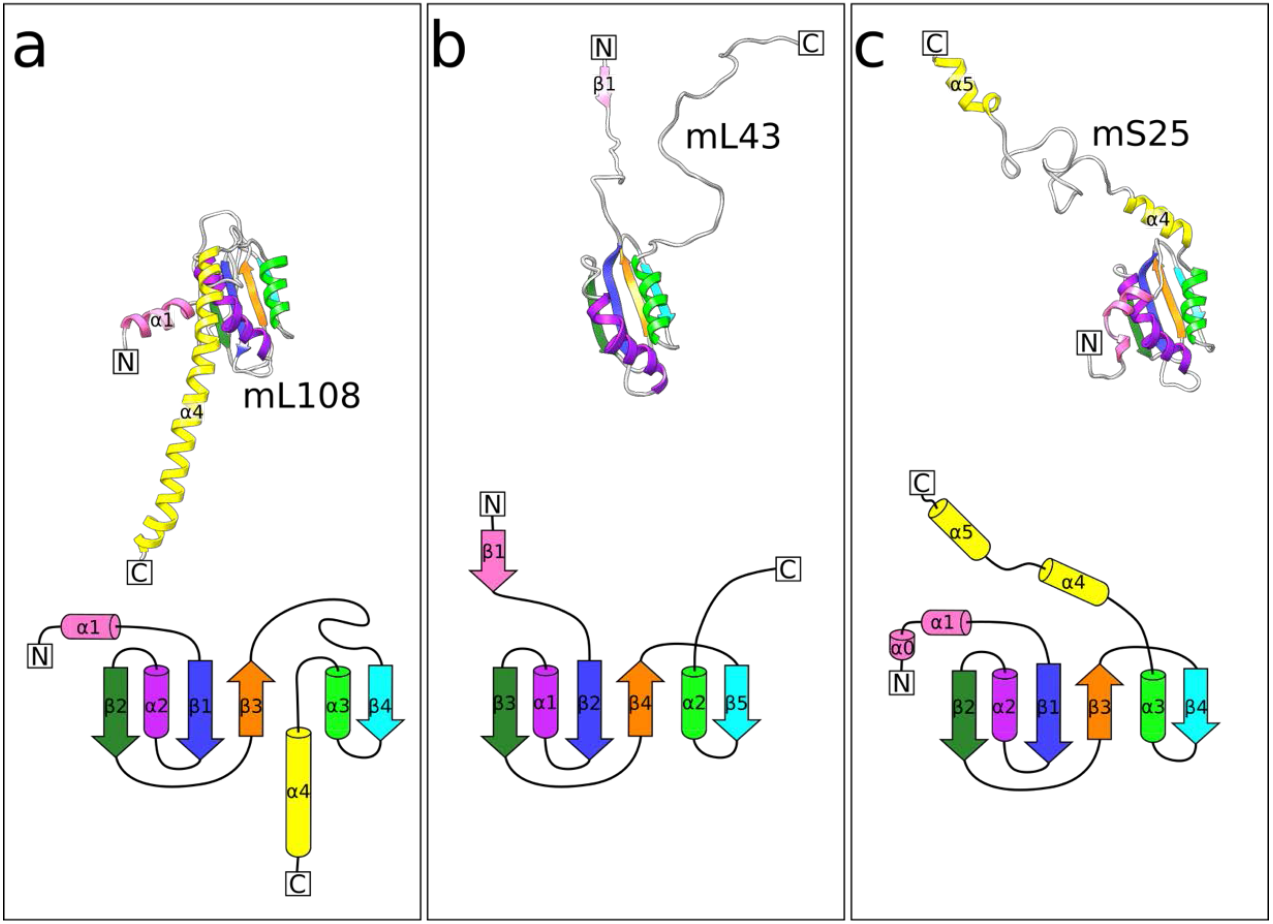
Topology of mL108, mL43, and mS25. Three dimensional models and topology of the homologous proteins with secondary structure elements colored, featuring their structural similarity. **a** mL108 of *N. crassa*. **b** mL43 of *N. crassa* and **c** mS25 of *Sus scrofa* (PDB ID: <u>5AJ4</u>).

**Supplementary Fig. 17:**
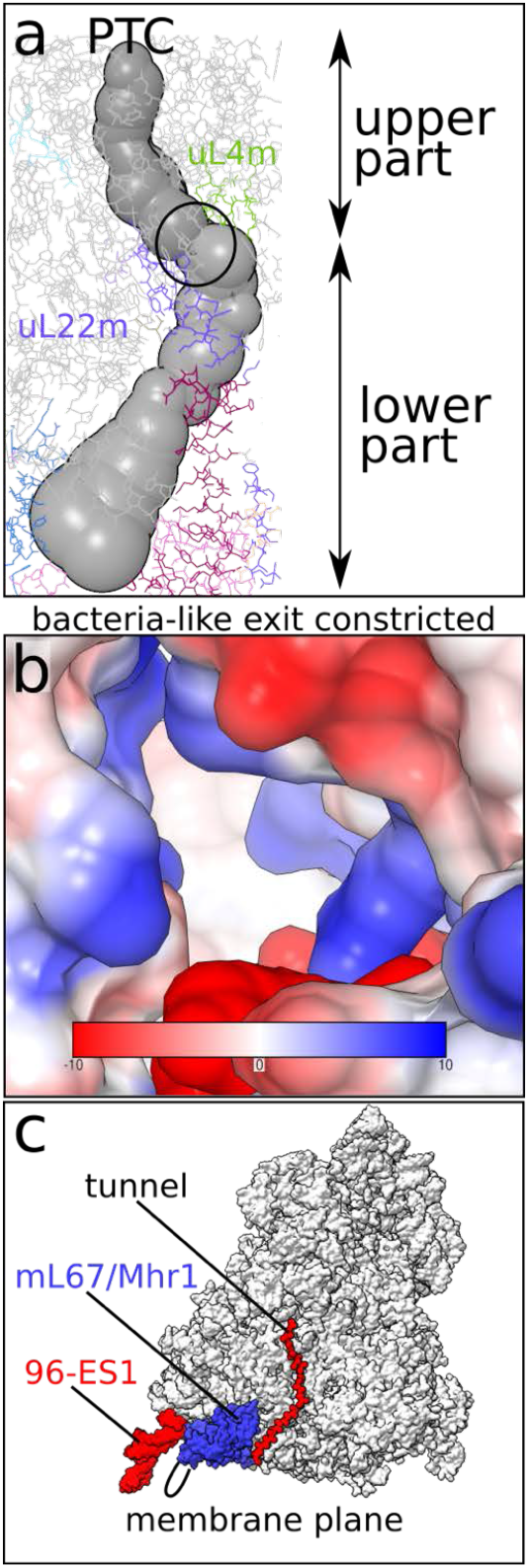
Analysis of potential tunnel exit sites. **a** The calculated tunnel showing the partition of the tunnel in to the upper and lower parts. The constriction site (circled) is surrounded by uL22m and uL4m. **b** The view of the bacteria-like exit site in surface representation colored by electrostatic Coulomb potential (kcal/mol*e) shows constriction with positively charged protein environment. **c** The polypeptide path in the *N. crassa* mitoribosome. The flexible elements 96-ES1 and mL67/Mhr1 extension around the exit are indicated.

**Supplementary Fig. 18:**
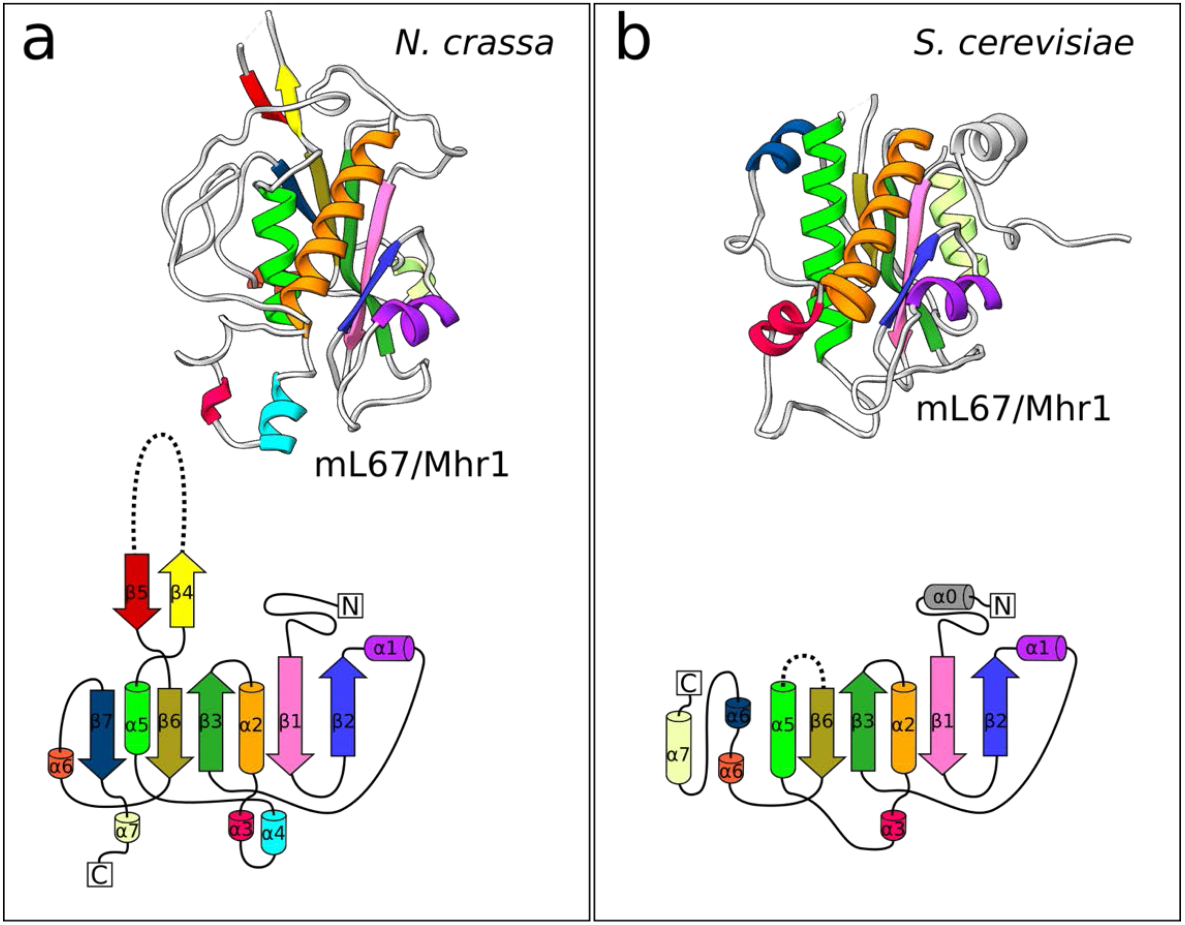
Structure and topology of mL67/Mhr1. **a** Three dimensional model and topology of *N. crassa* mL67/Mhr1 with secondary structure elements colored. The specific extension between β-strands 4 and 5 is indicated as dashed line. **b** Three dimensional model and topology of *S. cerevisiae* mL67/Mhr1 following the same color scheme in panel **a**.

**Supplementary Table 1.** Data and model statistics.

| Data collection | mtLSU | mtSSU | Monosome with E-site tRNA | Monosome with P-site tRNA | Monosome with tRNA in P/E state | Late stage intermediate mtLSU (Atp25) |
| --- | --- | --- | --- | --- | --- | --- |
| Microscope | Titan Krios | Titan Krios | Titan Krios | Titan Krios | Titan Krios | Titan Krios |
| Detector | K2 Summit | K2 Summit | K2 Summit | K2 Summit | K2 Summit | K2 Summit |
| Magnification | 130,000 | 130,000 | 130,000 | 130,000 | 130,000 | 130,000 |
| Voltage [kV] | 300 | 300 | 300 | 300 | 300 | 300 |
| Total electron dose [$e^-/\text{\AA}^2$] | 35 | 35 | 35 | 35 | 35 | 35 |
| Defocus range [$\mu\text{m}$] | -0.8 to -3.0 | -0.8 to -3.0 | -0.8 to -3.0 | -0.8 to -3.0 | -0.8 to -3.0 | -0.8 to -3.0 |
| Pixel size [$\text{\AA}$] | 1.06 | 1.06 | 1.06 | 1.06 | 1.06 | 1.06 |
| Final particles | 131,806 | 131,806 | 23,802 | 24,611 | 37,908 | 24,142 |
| Resolution [$\text{\AA}$] (overall/ LSU core / LSU CP / LSU L10 region / LSU L1 stalk / SSU head / SSU body / SSU tail) | 2.74/2.70/2.88/3.03/3.48/-/-/-t | 2.85/-/-/-/-/2.79/2.79/2.88 | 3.10/2.99/3.14/3.50/-/3.14/ 3.07/3.21 | 3.05/2.97/3.01/ 3.07/-/3.03/ 3.03/3.10 | 2.99/2.90/3.12/ 3.14/3.48/2.94/ 2.94/3.05 | 3.03/-/-/-/-/-/-/- |
| Map-sharpening $B$ factor [$\text{\AA}^2$] (overall/ LSU core / LSU CP / LSU L10 region / LSU L1 stalk / SSU head / SSU body / SSU tail) | -50.7/-48.3/-69.6/-74.4 /-66.7/-/-/- | -51.4/-/-/-/-/-48.3/-56.5/-64.7 | -41.2/-33.6/ -60.9/-55.2/-/ -50.4/-32.3/ -55.4 | -49.0/-36.8/ -46.0/-56.2/-/ -50.1/-50.1/ -57.0 | -45.8/-46.1/ -55.9/-55.9/ -66.7/-41.1/ -45.8/-61.0 | -40.3/-/-/-/-/-/-/- |
| **Model composition** |  |  |  |  |  |  |
| Total atoms (all atoms/ hydrogen) | 223,408/97,527 | 167,560/76,668 | 390,797/173,689 | 392,419/174,345 | 394,205/175,280 | 207,611/91,129 |
| Chains (RNA/ protein) | 1/44 | 1/35 | 3/78 | 4/79 | 4/80 | 1/42 |
| RNA residues | 2820 | 1435 | 4334 | 4362 | 4357 | 2529 |
| Protein residues | 8200 | 7528 | 15553 | 15604 | 15727 | 7802 |
| Metal ions ($\text{Mg}^{2+}$/ $\text{K}^+$/ $\text{Zn}^{2+}$) | 163/12/2 | 97/22/0 | 249/50/2 | 250/49/2 | 242/55/2 | 144/23/2 |
| Ligands (ATP/ NAD/ spermine) | 0/1/1 | 1/0/0 | 1/1/1 | 1/1/1 | 1/1/1 | 0/1/1 |
| **Refinement** |  |  |  |  |  |  |
| Model to map CC ($\text{CC}_{\text{mask}}$/ $\text{CC}_{\text{box}}$/ $\text{CC}_{\text{peaks}}$/ $\text{CC}_{\text{volume}}$) | 0.86/0.76/0.73/0.84 | 0.86/0.73/0.67/0.85 | 0.81/0.77/0.75/0.80 | 0.83/0.78/0.77/0.82 | 0.84/0.79/0.78/0.83 | 0.81/0.76/0.74/0.79 |
| Average $B$ factor [$\text{\AA}^2$] (protein/RNA/ metal ion and ligand/) | 42.56/39.71/23.92 | 36.80/36.55/31.32 | 42.75/53.92/29.43 | 21.37/21.50/11.39 | 27.67/29.15/18.25 | 17.73/25.77/15.02 |
| RMSD bond lengths [$\text{\AA}$] | 0.002 | 0.002 | 0.002 | 0.002 | 0.002 | 0.002 |
| RMSD bond angles [$^\circ$] | 0.382 | 0.357 | 0.373 | 0.354 | 0.362 | 0.402 |
| **Validation by MolProbity** |  |  |  |  |  |  |
| Clash score | 1.57 | 2.48 | 1.86 | 1.75 | 1.69 | 1.61 |
| Rotamer outliers [%] | 0.23 | 0.14 | 0.69 | 0.41 | 0.71 | 0.74 |
| Ramachandran plot [%] (Favored/<br>allowed/ disallowed) | 98.31/1.69/0.00 | 98.40/1.60/0.00 | 98.21/1.76/0.03 | 98.44/1.53/0.03 | 98.20/1.79/0.01 | 98.44/1.56/0.00 |
| **EMDB ID** (overall/ LSU core /<br>LSU CP / LSU L10 region / LSU<br>L1 stalk / SSU head / SSU body /<br>SSU tail) | 10973/10974/<br>10976/10975/-/-/-/- | 10958/-/-/-/-/<br>10961/10962/ 10963 | 10978/10979/10981/<br>10980/-/10982/<br>10984/10983 | 10985/10986/10989/<br>10988/-/10990/<br>10991/10992 | 10965/10966/10968/<br>10967/10971/10969/<br>10970/10972 | 10977/-/-/-/-/-/-/- |
| **PDB ID** | 6YWS | 6YW5 | 6YWX | 6YWY | 6YWE | 6YWV |

**Supplementary Table 2.**
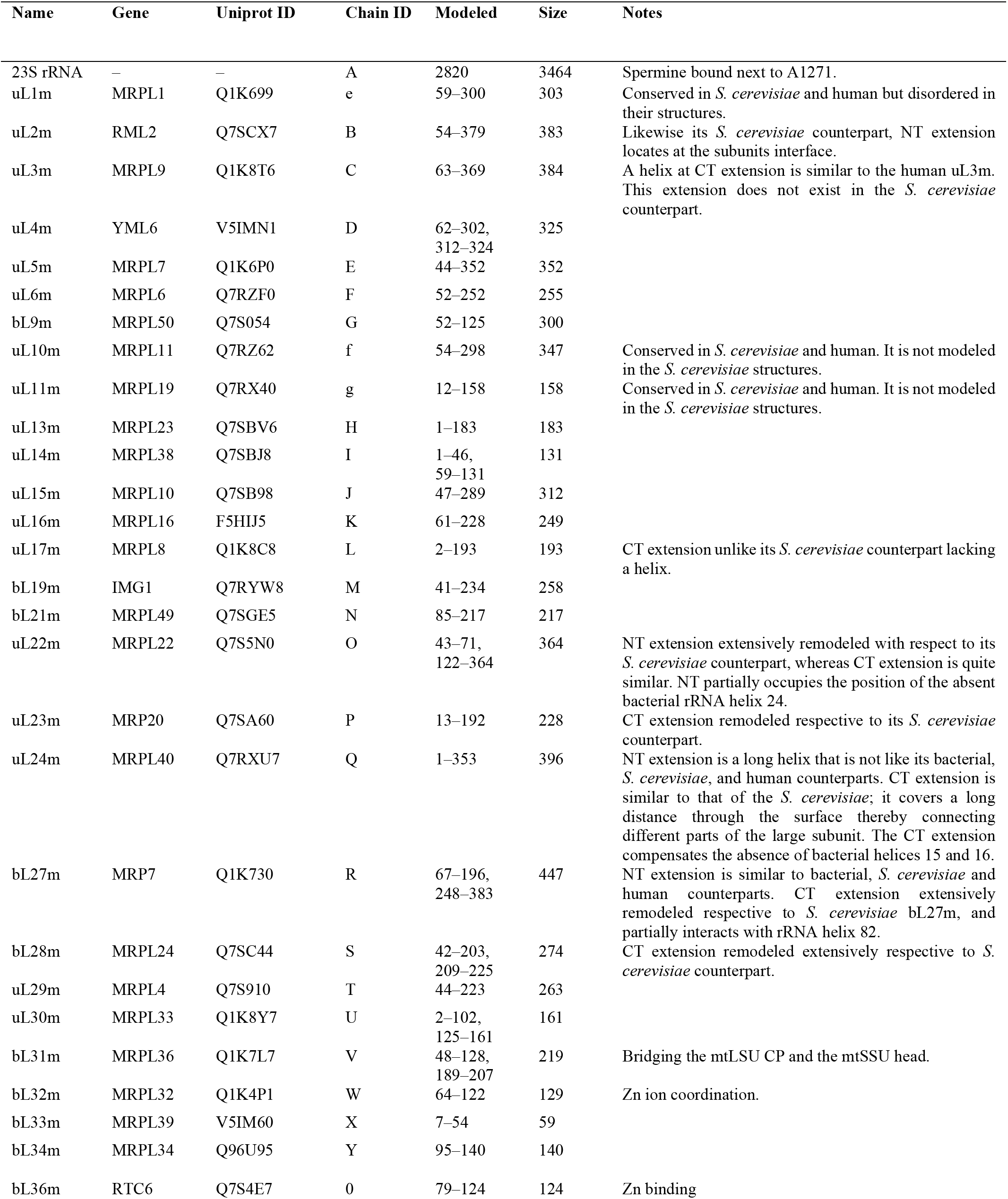
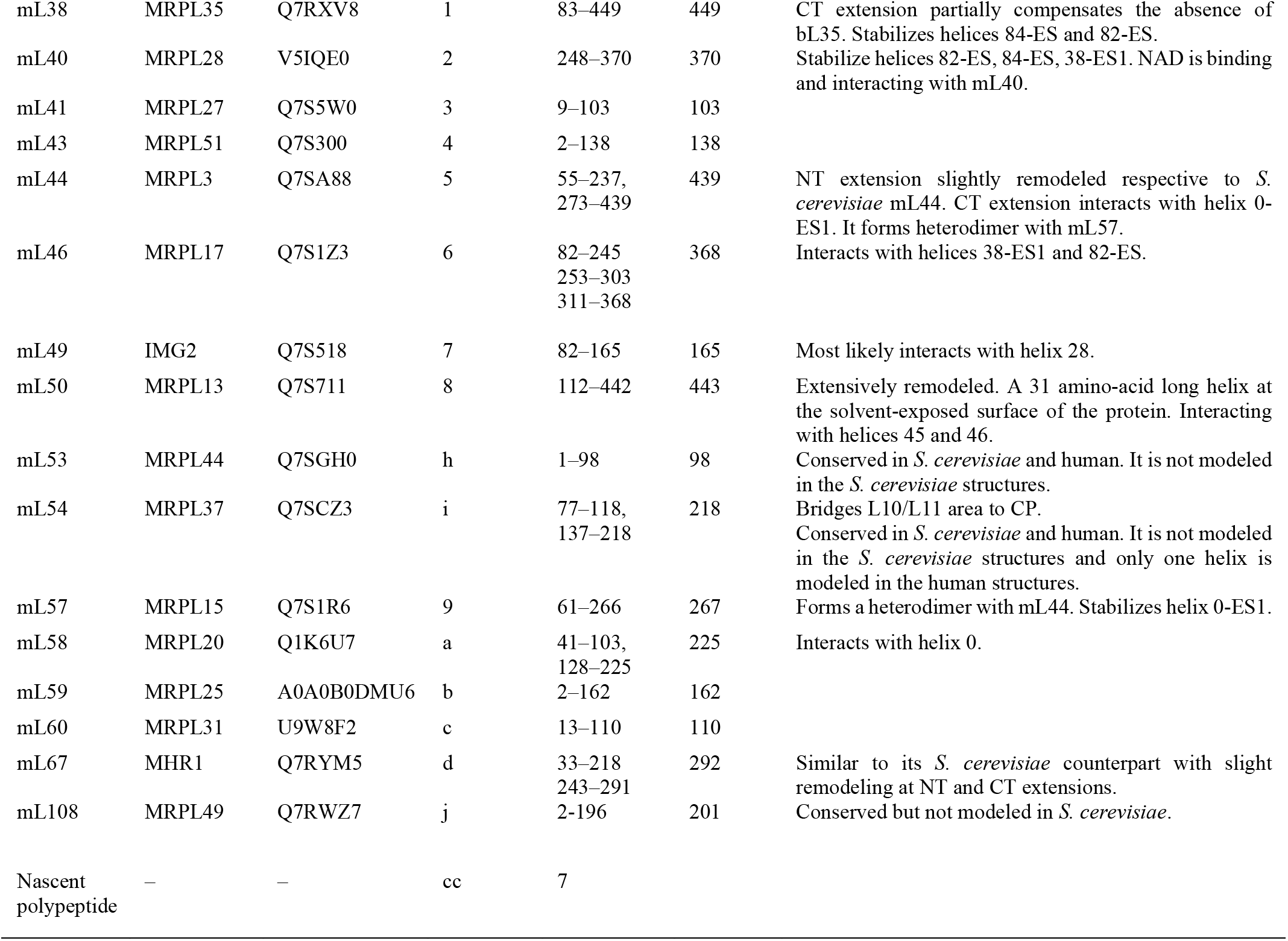
List of RNA and proteins from mtLSU.

**Supplementary Table 3.** List of RNA and proteins from mtSSU.

| MRP | Gene | Uniprot ID | Chain ID | Modeled | Size | Notes |
| --- | --- | --- | --- | --- | --- | --- |
| 16S rRNA | — | — | aa | 1435 | 1864 |  |
| bS1m | MRP51 | A7UWX2 | AA | 5–124,<br>132–213,<br>229–269,<br>288–337,<br>341–358,<br>368–430 | 470 |  |
| uS2m | MRP4 | V5ILE0 | BB | 107–396 | 428 |  |
| uS3m | VAR1 | P23351 | CC | 17–171,<br>193–369,<br>403–508 | 508 |  |
| uS4m | NAM9 | Q7SA90 | DD | 1–215,<br>379–453 | 453 |  |
| uS5m | MRPS5 | Q1K548 | EE | 56–180,<br>236–477 | 477 | Has a unique insertion to form a unique second beak. |
| bS6m | MRP17 | Q7SB95 | FF | 1–117 | 117 |  |
| uS7m | RSM7 | Q7S6M9 | GG | 87–309 | 309 |  |
| uS8m | MRPS8 | Q7SHF3 | HH | 2–161 | 161 |  |
| uS9m | MRPS9 | Q7S7R6 | II | 69–315 | 315 |  |
| uS10m | RSM10 | Q7RYL4 | JJ | 81–268 | 268 |  |
| uS11m | MRPS18 | Q7SGU0 | KK | 253–376 | 376 |  |
| uS12m | MRPS12 | Q7S9I4 | LL | 45–172 | 174 |  |
| uS13m | SWS2 | Q7S2C2 | MM | 2–119 | 119 |  |
| uS14m | MRP2 | Q7SF85 | NN | 2–113 | 113 |  |
| uS15m | MRPS28 | Q1K5G1 | OO | 38–106,<br>114–320 | 320 |  |
| bS16m | MRPS16 | P08580 | PP | 2–99 | 107 |  |
| uS17m | MRPS17 | Q7S4E0 | QQ | 6–163 | 165 | A unique C-terminal helix, which binds the IF<sub>1</sub> dimer. |
| bS18m | RSM18 | Q1K8E0 | RR | 108–241 | 256 |  |
| uS19m | RSM19 | Q1K8V2 | SS | 9–89 | 91 |  |
| bS21m | MRP21 | Q7SAJ1 | TT | 149–236 | 236 |  |
| mS23 | RSM25 | F8MRK5 | UU | 2–128,<br>142–238 | 240 |  |
| mS26 | PET123 | Q7SHR9 | VV | 47–257,<br>262–287,<br>293–316 | 316 |  |
| mS27 | MRP13 | Q7RYW7 | WW | 40–270,<br>275–396 | 396 | Corresponds to mS27 in human. *S. cerevisiae* mS44 and the unknown B (PDB ID: 5MRC) are one connected protein, which corresponds to human and *N. crassa* mS27. |
| mS29 | RSM23<br>/DAP3 | Q7SD06 | XX | 57–66,<br>72–469 | 469 | DAP3 death associated protein 3. ATP and Mg ion are bound. |
| mS33 | RSM27 | Q1K5R0 | YY | 2–100 | 108 |  |
| mS35 | RSM24 | Q1K5Z0 | ZZ | 42–353 | 382 | Has a unique N-terminal domain to form a unique beak. |
| mS37 | MRP10 | Q7S4Y4 | 11 | 2–89 | 90 |  |
| mS38 | COX24 | Q7SHR6 | 22 | 312–344 | 344 |  |
| mS41 | FYV4 | Q1K6Q3 | 33 | 43–235 | 236 |  |
| mS42 | RSM27<br>/MRP1 | Q7S2H6 | 44, 55 | 43–302 | 310 | Homolog of Fe superoxide dismutase. Lost the catalytic Fe ion binding side. Two chains form a homodimer, in contrast to the mS42-mS43 heterodimer in *S. cerevisiae*. |
| mS45 | MRPS35 | Q7SHB2 | 66 | 66–348 | 348 |  |
| mS46 | RSM28 | Q7SG49 | 77 | 223–401 | 414 |  |
| mS47 | MRP5<br>/EHD3 | Q1K7A4 | 88 | 42–508 | 508 | 3-hydroxyisobutyryl-CoA hydrolase. Probably an active enzyme because its active site is conserved. |
| ATP-IF<sub>1</sub> | INH1 | V5IRA3 | 00, 99 | 39–86 | 95 | ATP synthase inhibitor 1. Forms a homodimer located in the mtSSU tail region. |
| tRNA | – | – | bb | 73 |  | P, E or P/E tRNA state. The density is an average of different tRNA species. |
| mRNA | – | – | ee | 11 |  | mRNA in the P and P/E state. The density is an average of different mRNAs. |

## References

1 Brown, A. et al. Structure of the large ribosomal subunit from human mitochondria. Science 346, 718–722, doi:10.1126/science.1258026 (2014).

2 Amunts, A., Brown, A., Toots, J., Scheres, S. H. W. & Ramakrishnan, V. Ribosome. The structure of the human mitochondrial ribosome. Science 348, 95–98, doi:10.1126/science.aaa1193 (2015).

3 Greber, B. J. et al. Architecture of the large subunit of the mammalian mitochondrial ribosome. Nature 505, 515–519, doi:10.1038/nature12890 (2014).

4 Greber, B. J. et al. Ribosome. The complete structure of the 55S mammalian mitochondrial ribosome. Science 348, 303–308, doi:10.1126/science.aaa3872 (2015).

5 Amunts, A. et al. Structure of the yeast mitochondrial large ribosomal subunit. Science 343, 1485–1489, doi:10.1126/science.1249410 (2014).

6 Desai, N., Brown, A., Amunts, A. & Ramakrishnan, V. The structure of the yeast mitochondrial ribosome. Science 355, 528–531, doi:10.1126/science.aal2415 (2017).

7 van der Sluis, E. O. et al. Parallel Structural Evolution of Mitochondrial Ribosomes and OXPHOS Complexes. Genome Biol Evol 7, 1235–1251, doi:10.1093/gbe/evv061 (2015).

8 Kuntzel, H. & Noll, H. Mitochondrial and cytoplasmic polysomes from Neurospora crassa. Nature 215, 1340–1345, doi:10.1038/2151340a0 (1967).

9 Kuntzel, H. Mitochondrial and cytoplasmic ribosomes from Neurospora crassa: characterization of their subunits. J Mol Biol 40, 315–320, doi:10.1016/0022-2836(69)90481-1 (1969).

10 Neupert, W., Sebald, W., Schwab, A. J., Massinger, P. & Bucher, T. Incorporation in vivo of 14C-labelled amino acids into the proteins of mitochondrial ribosomes from Neurospora crassa sensitive to cycloheximide and insensitive to Chloramphenicol. Eur J Biochem 10, 589–591, doi:10.1111/j.1432-1033.1969.tb00730.x (1969).

11 Nargang, F. E., Preuss, M., Neupert, W. & Herrmann, J. M. The Oxa1 protein forms a homooligomeric complex and is an essential part of the mitochondrial export translocase in Neurospora crassa. J Biol Chem 277, 12846–12853, doi:10.1074/jbc.M112099200 (2002).

12 Brown, A. et al. Structures of the human mitochondrial ribosome in native states of assembly. Nat Struct Mol Biol 24, 866–869, doi:10.1038/nsmb.3464 (2017).

13 Kim, H. J., Maiti, P. & Barrientos, A. Mitochondrial ribosomes in cancer. Semin Cancer Biol 47, 67–81, doi:10.1016/j.semcancer.2017.04.004 (2017).

14 Robinson, G. C. et al. The structure of F(1)-ATPase from Saccharomyces cerevisiae inhibited by its regulatory protein IF(1). Open Biol 3, 120164, doi:10.1098/rsob.120164 (2013).

15 Cabezon, E., Montgomery, M. G., Leslie, A. G. & Walker, J. E. The structure of bovine F1-ATPase in complex with its regulatory protein IF1. Nat Struct Biol 10, 744–750, doi:10.1038/nsb966 (2003).

16 Cabezon, E., Runswick, M. J., Leslie, A. G. & Walker, J. E. The structure of bovine IF(1), the regulatory subunit of mitochondrial F-ATPase. EMBO J 20, 6990–6996, doi:10.1093/emboj/20.24.6990 (2001).

17 Petrov, A. S. et al. Structural Patching Fosters Divergence of Mitochondrial Ribosomes. Mol Biol Evol 36, 207–219, doi:10.1093/molbev/msy221 (2019).

18 Qu, X., Lancaster, L., Noller, H. F., Bustamante, C. & Tinoco, I., Jr. Ribosomal protein S1 unwinds double-stranded RNA in multiple steps. Proc Natl Acad Sci U S A 109, 14458–14463, doi:10.1073/pnas.1208950109 (2012).

19 Komoda, T. et al. The A-site finger in 23 S rRNA acts as a functional attenuator for translocation. J Biol Chem 281, 32303–32309, doi:10.1074/jbc.M607058200 (2006).

20 Aibara, S., Singh, V., Modelska, A. & Amunts, A. Structural basis of mitochondrial translation. Elife 9, doi:10.7554/eLife.58362 (2020).

21 Tobiasson, V. & Amunts, A. Ciliate mitoribosome illuminates evolutionary steps of mitochondrial translation. Elife 9, doi:10.7554/eLife.59264 (2020).

22 Kummer, E. & Ban, N. Structural insights into mammalian mitochondrial translation elongation catalyzed by mtEFG1. EMBO J 39, e104820, doi:10.15252/embj.2020104820 (2020).

23 Dao Duc, K., Batra, S. S., Bhattacharya, N., Cate, J. H. D. & Song, Y. S. Differences in the path to exit the ribosome across the three domains of life. Nucleic Acids Res 47, 4198–4210, doi:10.1093/nar/gkz106 (2019).

24 Ling, F. & Shibata, T. Recombination-dependent mtDNA partitioning: in vivo role of Mhr1p to promote pairing of homologous DNA. EMBO J 21, 4730–4740, doi:10.1093/emboj/cdf466 (2002).

25 Ling, F. & Shibata, T. Mhr1p-dependent concatemeric mitochondrial DNA formation for generating yeast mitochondrial homoplasmic cells. Mol Biol Cell 15, 310–322, doi:10.1091/mbc.e03-07-0508 (2004).

26 Prasai, K., Robinson, L. C., Scott, R. S., Tatchell, K. & Harrison, L. Evidence for double-strand break mediated mitochondrial DNA replication in Saccharomyces cerevisiae. Nucleic Acids Res 45, 7760–7773, doi:10.1093/nar/gkx443 (2017).

27 Woellhaf, M. W., Sommer, F., Schroda, M. & Herrmann, J. M. Proteomic profiling of the mitochondrial ribosome identifies Atp25 as a composite mitochondrial precursor protein. Mol Biol Cell 27, 3031–3039, doi:10.1091/mbc.E16-07-0513 (2016).

28 Kudva, R. et al. The shape of the bacterial ribosome exit tunnel affects cotranslational protein folding. Elife 7, doi:10.7554/eLife.36326 (2018).

29 Stoltzfus, A. Constructive neutral evolution: exploring evolutionary theory’s curious disconnect. Biol Direct 7, 35, doi:10.1186/1745-6150-7-35 (2012).

30 Hauser, R. et al. RsfA (YbeB) proteins are conserved ribosomal silencing factors. PLoS Genet 8, e1002815, doi:10.1371/journal.pgen.1002815 (2012).

31 Li, X. et al. Structure of Ribosomal Silencing Factor Bound to Mycobacterium tuberculosis Ribosome. Structure 23, 2387, doi:10.1016/j.str.2015.11.002 (2015).

32 Rorbach, J., Gammage, P. A. & Minczuk, M. C7orf30 is necessary for biogenesis of the large subunit of the mitochondrial ribosome. Nucleic Acids Res 40, 4097–4109, doi:10.1093/nar/gkr1282 (2012).

33 Zeng, X., Barros, M. H., Shulman, T. & Tzagoloff, A. ATP25, a new nuclear gene of Saccharomyces cerevisiae required for expression and assembly of the Atp9p subunit of mitochondrial ATPase. Mol Biol Cell 19, 1366–1377, doi:10.1091/mbc.E07-08-0746 (2008).

34 Weis, F. et al. Mechanism of eIF6 release from the nascent 60S ribosomal subunit. Nat Struct Mol Biol 22, 914–919, doi:10.1038/nsmb.3112 (2015).

35 Bason, J. V., Montgomery, M. G., Leslie, A. G. & Walker, J. E. Pathway of binding of the intrinsically disordered mitochondrial inhibitor protein to F1-ATPase. Proc Natl Acad Sci U S A 111, 11305–11310, doi:10.1073/pnas.1411560111 (2014).

36 Funes, S., Nargang, F. E., Neupert, W. & Herrmann, J. M. The Oxa2 protein of Neurospora crassa plays a critical role in the biogenesis of cytochrome oxidase and defines a ubiquitous subbranch of the Oxa1/YidC/Alb3 protein family. Mol Biol Cell 15, 1853–1861, doi:10.1091/mbc.e03-11-0789 (2004).

37 Vogel, H. J. A convenient growth medium for Neurospora crassa. Microbial Genet Bull. 13, 42–43 (1956).

38 Li, X. et al. Electron counting and beam-induced motion correction enable near-atomic-resolution single-particle cryo-EM. Nat Methods 10, 584–590, doi:10.1038/nmeth.2472 (2013).

39 Zhang, K. Gctf: Real-time CTF determination and correction. J Struct Biol 193, 1–12, doi:10.1016/j.jsb.2015.11.003 (2016).

40 de la Rosa-Trevin, J. M. et al. Scipion: A software framework toward integration, reproducibility and validation in 3D electron microscopy. J Struct Biol 195, 93–99, doi:10.1016/j.jsb.2016.04.010 (2016).

41 Kimanius, D., Forsberg, B. O., Scheres, S. H. & Lindahl, E. Accelerated cryo-EM structure determination with parallelisation using GPUs in RELION-2. Elife 5, doi:10.7554/eLife.18722 (2016).

42 Zivanov, J. et al. New tools for automated high-resolution cryo-EM structure determination in RELION-3. Elife 7, doi:10.7554/eLife.42166 (2018).

43 Zivanov, J., Nakane, T. & Scheres, S. H. W. A Bayesian approach to beam-induced motion correction in cryo-EM single-particle analysis. IUCrJ 6, 5–17, doi:10.1107/S205225251801463X (2019).

44 Emsley, P. & Cowtan, K. Coot: model-building tools for molecular graphics. Acta Crystallogr D Biol Crystallogr 60, 2126–2132, doi:10.1107/S0907444904019158 (2004).

45 Pettersen, E. F. et al. UCSF Chimera--a visualization system for exploratory research and analysis. J Comput Chem 25, 1605–1612, doi:10.1002/jcc.20084 (2004).

46 Waterhouse, A. et al. SWISS-MODEL: homology modelling of protein structures and complexes. Nucleic Acids Res 46, W296–W303, doi:10.1093/nar/gky427 (2018).

47 Altschul, S. F., Gish, W., Miller, W., Myers, E. W. & Lipman, D. J. Basic local alignment search tool. J Mol Biol 215, 403–410, doi:10.1016/S0022-2836(05)80360-2 (1990).

48 Ban, N. et al. A new system for naming ribosomal proteins. Curr Opin Struct Biol 24, 165–169, doi:10.1016/j.sbi.2014.01.002 (2014).

49 Liebschner, D. et al. Macromolecular structure determination using X-rays, neutrons and electrons: recent developments in Phenix. Acta Crystallogr D Struct Biol 75, 861–877, doi:10.1107/S2059798319011471 (2019).

50 Afonine, P. V. et al. Real-space refinement in PHENIX for cryo-EM and crystallography. Acta Crystallogr D Struct Biol 74, 531–544, doi:10.1107/S2059798318006551 (2018).

51 Williams, C. J. et al. MolProbity: More and better reference data for improved all-atom structure validation. Protein Sci 27, 293–315, doi:10.1002/pro.3330 (2018).

52 DeLano, W. L. Pymol: An open-source molecular graphics tool.. CCP4 Newsletter On Protein Crystallography 40, 82–92 (2002).

53 Goddard, T. D. et al. UCSF ChimeraX: Meeting modern challenges in visualization and analysis. Protein Sci 27, 14–25, doi:10.1002/pro.3235 (2018).

54 Pravda, L. et al. MOLEonline: a web-based tool for analyzing channels, tunnels and pores (2018 update). Nucleic Acids Res 46, W368–W373, doi:10.1093/nar/gky309 (2018).

## Supplementary References

1 Cabezon, E., Runswick, M. J., Leslie, A. G. & Walker, J. E. The structure of bovine IF(1), the regulatory subunit of mitochondrial F-ATPase. EMBO J 20, 6990–6996, doi:10.1093/emboj/20.24.6990 (2001).

2 Bason, J. V., Montgomery, M. G., Leslie, A. G. & Walker, J. E. Pathway of binding of the intrinsically disordered mitochondrial inhibitor protein to F1-ATPase. Proc Natl Acad Sci U S A 111, 11305–11310, doi:10.1073/pnas.1411560111 (2014).

